# Machine Learning Models for Predicting Multiple Myeloma Staging and MGUS Progression Using Gene Expression Data

**DOI:** 10.1101/2024.11.12.623149

**Authors:** Nestoras Karathanasis, George M. Spyrou

## Abstract

In this study, we developed and evaluated Machine Learning (ML) models aimed at predicting the stage of multiple myeloma (MM) and the progression of monoclonal gammopathy of undetermined significance (MGUS) to MM. Accurate staging of MM is critical for determining appropriate treatment strategies, and our models, employing algorithms such as ElasticNet, Random Forest, Boosting, and Support Vector Machines, demonstrated high efficacy in capturing the biological differences across disease stages. Among these, the ElasticNet model exhibited strong generalizability, achieving consistent multiclass AUC values across various datasets and data transformations.

Predicting MGUS progression to MM presents a significant challenge due to the scarcity of MGUS cases that have progressed. We employed a two-pronged approach to address this: developing models using a limited dataset containing progressing MGUS patients and training models on combined MGUS and MM datasets. The models achieved AUC values slightly above 0.8, particularly with ElasticNet, Boosting and Support Vector Machines, indicating their potential in stratifying MGUS patients by progression risk. This study is original in integrating MM data with MGUS cases to enhance the predictive accuracy of MGUS progression, offering a novel methodology with potential clinical applications in patient monitoring and early intervention.

Our feature selection and enrichment analyses further revealed that the identified genes are involved in key signaling pathways, including PI3K-Akt, MAPK, Wnt, and mTOR, all of which play crucial roles in MM pathogenesis. These findings align with established biological knowledge, suggest possible therapeutic targets and increase the explainability of our models.

## 1 Introduction

Multiple myeloma constitutes approximately 1% of all cancer cases and about 10% of hematologic malignancies ^1,2^. Annually, more than 32,000 new cases are diagnosed in the United States, with nearly 13,000 resulting in fatalities ^3^. The yearly age-adjusted incidence has remained steady for decades, hovering around 4 per 100,000 individuals ^4^. It shows a slight preference for men over women and is twice as prevalent among African Americans compared to Caucasians ^5^. The median age at diagnosis is typically around 65 years ^6^.

Nearly all multiple myeloma patients progress from an asymptomatic precursor stage known as monoclonal gammopathy of undetermined significance (MGUS) ^7,8^. MGUS is found in roughly 5% of individuals aged over 50, with a prevalence around twice as high among Blacks compared to Whites ^9–12^. MGUS transitions to multiple myeloma or related malignancies at a rate of 1% per year ^13,14^. As MGUS is asymptomatic, over 50% of those diagnosed with it have likely harboured the condition for over a decade before clinical diagnosis ^15^. In particular cases, an intermediate asymptomatic but more advanced pre-malignant stage, termed smoldering multiple myeloma (SMM), may be clinically recognisable ^16^. SMM progresses to multiple myeloma at a rate of approximately 10% per year within the first five years post-diagnosis, followed by 3% annually over the subsequent five years and 1.5% per year thereafter. This progression rate is influenced by the underlying cytogenetic profile, with patients harbouring specific translocations at a higher risk of progressing from MGUS or SMM to multiple myeloma ^17–19^.

Despite notable therapeutic advancements in recent years, multiple myeloma (MM) remains an uncurable disease. Enhanced insights into MM’s biology and pathogenesis have prompted a transformative shift in managing MM and its precursor states, monoclonal gammopathy of undetermined significance (MGUS) and smoldering multiple myeloma (SMM) ^20^. The conventional notion that MM treatment should only start upon the onset of symptoms has been challenged by the introduction of novel therapies characterised by both safety and efficacy. Clinical trials have underscored the significance of initiating treatment early in high-risk asymptomatic cases, demonstrating a marked delay in disease progression and improved progression-free survival outcomes for patients ^21,22^.

Yet, a critical challenge persists in identifying individuals with asymptomatic myeloma at the highest risk of progression, thereby maximising the benefits of early treatment strategies. While risk stratification models such as the Mayo Clinic model ^23^ and the Spanish model ^24^ have been valuable, they still possess notable limitations, particularly in the context of modern therapies. Studies have revealed that patients with high-risk cytogenetic MM, including del17p, t(4;14), or t(14;20), may achieve survival rates comparable to standard-risk patients through intensified treatment regimens involving a combination of proteasome inhibitors, immunomodulatory drugs, and autologous stem cell transplantation ^25^. Consequently, there is an urgent imperative to deepen our comprehension of the molecular mechanisms underpinning disease progression and refine risk stratification models for asymptomatic MM concurrently with endeavours to optimise early treatment strategies.

Over the past several years, there has been a notable increase in the utilization of machine learning (ML) algorithms and deep learning (DL) procedures for tumor detection. To manage cancer patients, these methods leverage diverse data sources such as proteomic, genomic, histopathological data, or images. These techniques have proven to be beneficial not only in the realm of solid tumors but also in the management of hematological malignancies. Recent reports on multiple myeloma have highlighted the significance of machine learning in diagnosing, prognosticating, response to treatment and evaluating therapeutic responses in hematological neoplasms ^26^. Currently, serum markers are employed to categorize MGUS patients into different clinical risk groups. However, no established molecular signature can reliably predict the progression of MGUS. To address this gap, Sun et al. ^27^ conducted a study utilizing gene expression profiling to stratify the risk of MGUS and devised a signature based on extensive samples with long-term follow-up. They analyzed microarrays of plasma cell mRNA from 334 MGUS patients with stable disease and 40 MGUS patients who progressed to multiple myeloma (MM) within a decade and identified a thirty-six-gene molecular signature indicative of MGUS risk.

The objectives of this study are: (1) develop machine-learning models capable of accurately predicting the stage of multiple myeloma (MM) based on microarray datasets. We utilized advanced algorithms to analyse molecular data to classify patients into different stages of the disease, thereby aiding clinicians in making more informed treatment decisions. (2) To investigate the feasibility of using machine-learning techniques to predict disease progression from monoclonal gammopathy of undetermined significance (MGUS) to multiple myeloma (MM). By leveraging microarray datasets containing gene expression profiles and clinical information from patients at the MGUS stage, the study aims to develop predictive models that identify individuals at high risk of progressing to MM. Models trained to distinguish MGUS from MM were tested for their effectiveness in identifying progressing MGUS cases ^27^, with results indicating similar or better performance to models explicitly trained for this task. This proactive approach aims to enable early intervention strategies and improve patient outcomes by potentially delaying or preventing disease progression.

## 2 Method

### 2.1 Source of Microarray Datasets and Description of Data Variables and Features

We downloaded seven microarray datasets, two from ArrayExpress and five from the Gene Expression Omnibus. In all cases, the samples were CD-138+ bone marrow plasma cells from patients with different stages of multiple myeloma (MGUS, MM) and Healthy. The datasets come from four different platforms (A-AFFY-33, A-AFFY-44, GPL96, GPL570) and contain different numbers of patients in total and per stage, see Table 1.

**Table 1.** The number of samples per dataset and disease stage. The table is sorted by the total number of samples. The empty cells correspond to a zero number of samples.

| Platform | Dataset | Normal | MGUS | progressing MGUS | MM | Total number of samples |
| --- | --- | --- | --- | --- | --- | --- |
| GPL96 | GSE2113 |  | 7 |  | 39 | 46 |
| A-AFFY-33<br>A-AFFY-34 | EMTAB316 |  | 7 |  | 65 | 72 |
| GPL570 | GSE5900 | 22 | 44 |  |  | 66 |
| GPL96 | GSE6477 | 15 | 22 |  | 73 | 110 |
| GPL96 | GSE13591 | 5 | 11 |  | 133 | 149 |
| A-AFFY-44 | EMTAB317 |  | 23 |  | 226 | 249 |
| GPL570 | GSE235356 |  | 319 | 39 |  | 358 |
| **Total** | **7** | **42** | **433** | **39** | **536** | **1050** |

For each dataset, we downloaded the raw .cel files. We calculated the expression matrix using the “Robust Multi-Array Average” expression measure via the rma() function of the affy or oligo R packages, depending on the requirements of each dataset, with background correction. At this step of the analysis, data was not normalized. Datasets from different platforms have different numbers of probes. GLP96, A-AFFY-33/A-AFFY-34 contain ∼22.000 probes, whereas GLP570 and A-AFFY-44 contain ∼55.000. We retained only the 22.277 shared probes across all datasets. Also, datasets contained samples corresponding to disease stages outside of the scope of this study (for example, SMM, relapse MM, PCL, and HUVEC) which we removed from our analysis.

### 2.2 Data Cleaning and Pre-processing Techniques

We employed several data transformation and normalization techniques to prepare our datasets for analysis, (refer to Figure 1):

**Figure 1.**
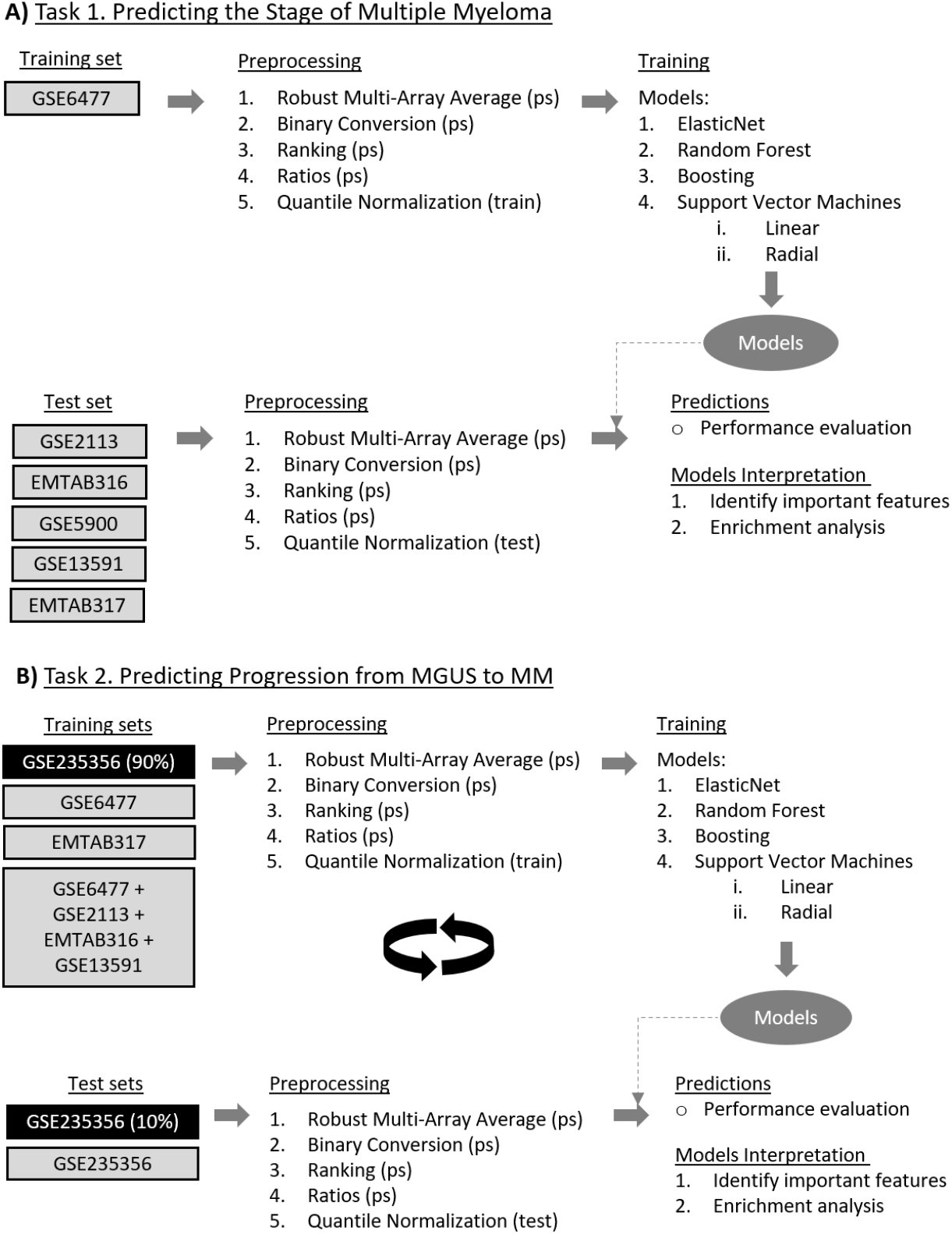
Flowchart of the analysis. **A)** The flowchart illustrates the process used for predicting the stage of Multiple Myeloma. The method encompasses multiple steps: data preprocessing, model training, and performance evaluation, applied across various datasets. Preprocessing includes several data transformations and the training phase incorporates a variety of machine learning models. After predictions, the model’s key features were interpreted through enrichment analyses. In the figure, (ps) indicates per-sample preprocessing, (train) denotes that normalization was applied to training samples, and (test) refers to applying the parameters learned from training to the test set. **B)** The flowchart outlines the process used for predicting the progression of MGUS to MM using machine learning techniques. The method involves preprocessing, model training, and performance evaluation using different datasets similar to A. The boxes with a black background indicate the use of the GSE235356 dataset for training and testing in a 10-fold nested cross-validation fashion. In contrast, grey background boxes represent training on various datasets and testing on the GSE235356 dataset.

#### Robust Multi-Array Average (rma)

We utilized rma function with data background correction, which is implemented in affy or oligo R packages, depending on the requirements of each dataset.

#### Binary Conversion

Expression values from rma were converted to binary (0-1) using two quantile thresholds, 0 (binary_0) and 0.5 (binary_0.5) per sample. Values exceeding the quantile threshold were set to 1, while those equal to or below the threshold were set to 0. In the case of binary_0, all values except the minimum were set to 1, and the minimum value was set to 0. Binary_0 was used as a negative control, where we expected the machine learning algorithms to perform as random classifiers, offering a baseline for performance comparison.

#### Ranking (ranking)

Expression values were ranked from 0 to 1, with the highest value assigned a rank of 1. This ranking system provided a relative measure of gene expression levels within each sample.

#### Ratios (ratio)

We selected only healthy samples from the GSE6477 dataset, which served solely as a training set. We calculated the ratios performing the following steps. First, we used the ranks from the ranking transformation, and calculated the standard deviation of each probe. We kept 210 probes with the lowest standard deviation. This number was chosen to minimize feature combinations, as the total number of combinations when selecting two genes each time was 21945 features—close to the total number of features from the other pre-processing approaches. Then we calculated all possible ratios of these probes.

#### Quantile Normalization (qnorm)

Quantile normalization was applied in a train-test fashion using the preprocess R package. The training set underwent quantile normalization, and the parameters learned from this process were then applied to the test set. This approach ensured consistency in data distribution between the training and test datasets.

### 2.3 Overview of Machine Learning (ML) Algorithms

We evaluated the following parametric and non-parametric methods, (see Figure 1).

***ElasticNet*** (glmnet) is a parametric method that fits generalized linear and similar models via penalized maximum likelihood^28^. We employed its implementation in the glmnet package in R. ElasticNet’s advantage is that it is the most interpretable ML method ^29^ among these mentioned here.

***Random Forest*** (rf) is a non-parametric tree-based method. We utilized its implementation in the randomForest R package. RF is somewhat interpretable as it provides information on which features are more important for the model by calculating variable importance scores ^29^.

***Boosting*** (gbm) is a non-parametric method. We used gradient boosting machines implemented in the gbm package in R. Like RF, boosting is somewhat interpretable and provides the most important features ^30^.

***Support Vector Machines (SVM)*** is a non-parametric method that has the advantage of projecting the data to a different feature space ^29^. However, even though SVMs can produce very accurate models, they lack interpretability. We used the implementation of SVMs in the e1071^31^ R package to fit an SVM with the linear kernel (svmLinear2) and the implementation in the kernlab ^32^ R package to fit an SVM with the radial kernel (svmRadial).

### 2.4 Models Training and Interpretation

We utilised the caret R package, which stands for classification and regression training ^33^, to train, optimise, and test our models, (see Figure 1). In order to optimize the model’s hyperparameters, we employed a ten-fold cross-validation repeated ten times. As the performance metric to determine the best model, we used the multiclass area under the ROC curve (multiclass_auc) for multiclass problems (see task 1 below) and the area under the ROC curve (AUC) for two-class problems (see task 2 below). For all models, except svmRadial, we tuned our models in a set of ten hyperparameters by setting caret’s tuneLength argument to ten. For the svmRadial model, we used the sigest() function from the kernlab R package to calculate the range of the sigma hyperparameter. The cost hyperparameter was set to the following values: 0.25, 0.50, 2, 4, 8, 16, 32, 64, 128, 256, 512, 1024. In all cases, the data were centred and scaled.

To interpret our models, we calculated the importance of each feature by utilizing the varImp() function from the caret R package. We then performed enrichment analysis for GO biological processes, KEGG and Reactome pathways, and disease ontology semantics using the clusterProfiler R package ^34^.Last, we filtered the results with the following terms related to multiple myeloma, MAPK, RAS, RAF, MEK, ERK, ERK1, ERK2, PI3K, AKT, NF-KB, Jak-STAT, Wnt, Hedgehog, TNFa, mTOR, multiple myeloma, myeloid, leukemia, myeloma, Plasmacytoma, Amyloidosis, Chronic Lymphocytic Leukemia, Heavy Chain Disease, Lymphoma ^35,36^.

## 3 Results

### 3.1 Task 1 - Predicting the Stage of Multiple Myeloma

#### 3.1.1 Model Development for Disease Staging

We trained our models using the GSE6477 microarray dataset ^37^. This dataset comprises 162 samples representing various stages of myeloma. Specifically, it includes 15 samples classified as Normal, 21 as MGUS (Monoclonal Gammopathy of Undetermined Significance), 23 as SMM (Smoldering Multiple Myeloma), 75 as MM (newly diagnosed myeloma), and 28 as RMM (relapsed myeloma samples). We focused on 110 samples after excluding the SMM and RMM categories. We trained our models to separate the three classes: Normal, MGUS and MM. We used all other datasets (see Table 1) only for testing.

#### 3.1.2 Evaluation of Model Performance

During training, all models consistently achieved a multiclass_auc with a cross-validation median ranging from 0.9 to 1 across various data transformations and machine-learning methods (refer to Supplementary Figure 1). For the binary_0 transformation, where all expressions except from the lowest were set to 1, the median training performance of all models was around 0.5, that corresponds to random classifier, as expected. Subsequently, we evaluated the multiclass_auc for all test datasets using all models and data transformations (see Figure 2 and Supplementary Figure 2).

**Figure 2.**
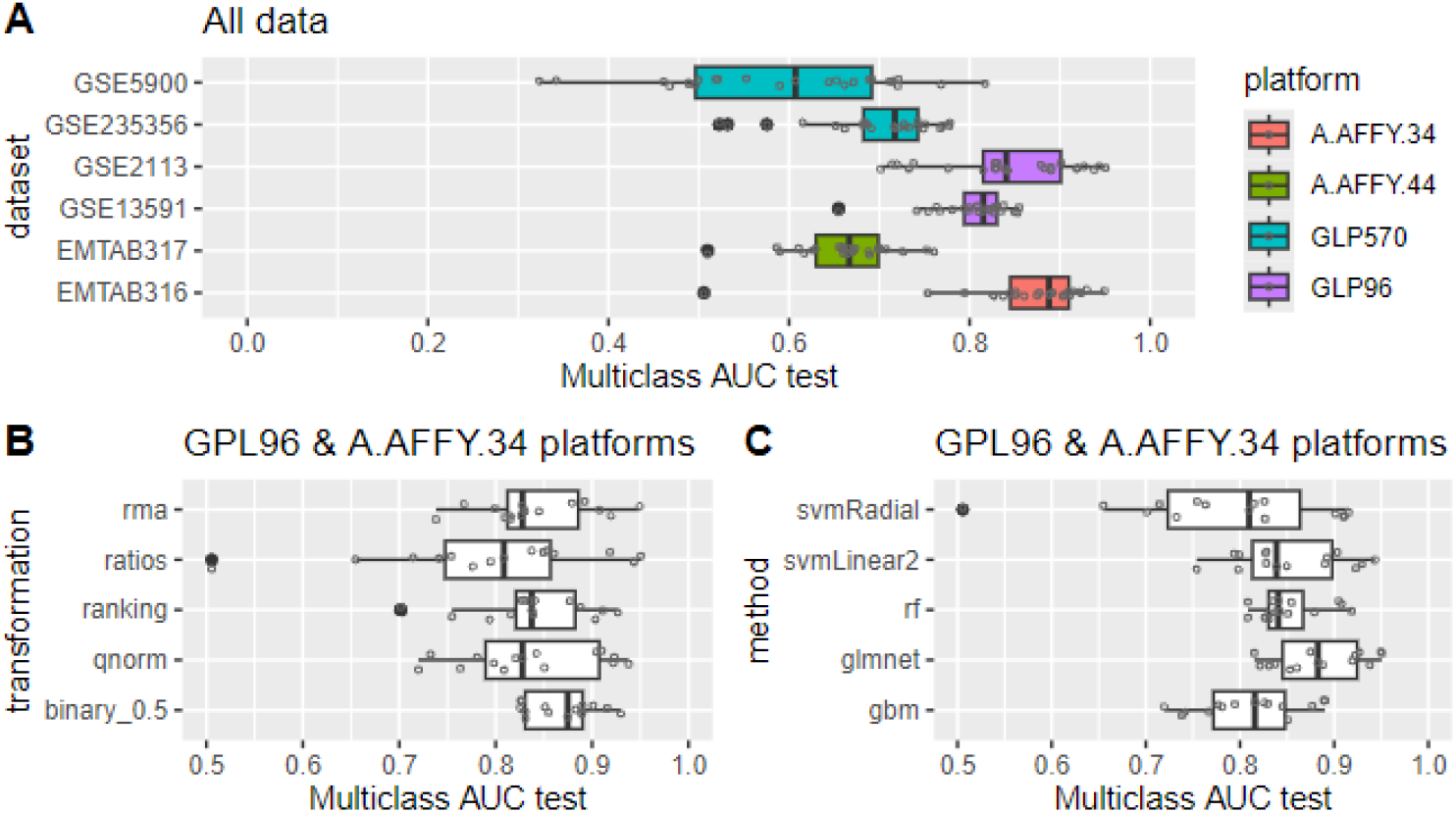
Models multiclass auc in the external validation sets. A) The performance to the external dataset used across all data transformations and machine learning algorithms. B) The relation of performance to the data transformations across datasets generated in GLP96 or A.AFFY.34 platforms and all machine learning algorithms. C) The relation of performance to the machine learning algorithms across datasets generated in GLP96 or A.AFFY.34 platforms and all data transformations.

Focusing on the platform of origin for the data, we noted that datasets (GSE13591, GSE2113) originating from the same platform (GPL96) as the training set exhibited similar multiclass_auc scores as seen during training, (see Figure 2A). We observed a slight decline in performance, approximately 0.1 (see Supplementary Figure 3). EMTAB316 originated from A-AFFY-34, which is very close to GPL96, showed similar multiclass_auc scores with training. In contrast, for datasets generated using different platforms (EMTAB317 from A-AFFY-44 and GSE5900, GSE235356 from GPL570), our models experienced a more significant decrease in performance. This suggests that performance variability across datasets may be attributed to differences in the platforms used for data generation. Regarding GSE235356, it’s important to note that this dataset includes only MGUS and progressing MGUS samples, which could contribute to the observed decline in performance.

**Figure 3.**
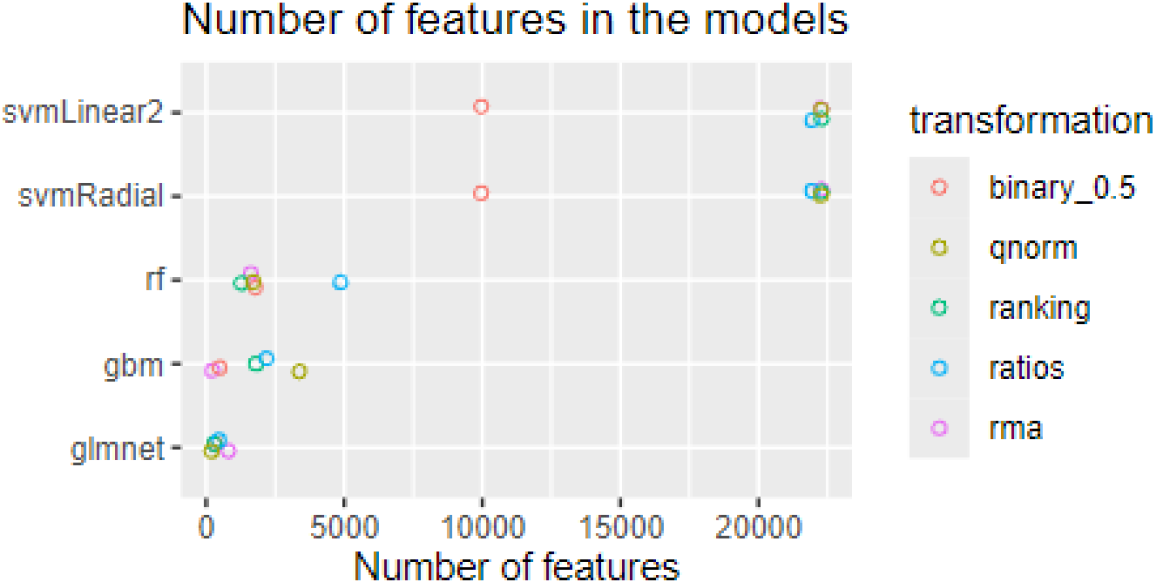
The number of features utilized by each model across different data transformations. The plot shows the variation in feature selection for each model, highlighting the range of features used in the analysis.

For the subsequent phases of our analysis, we focused on datasets generated from the GPL96 and A-AFFY-34 platforms. Concerning data transformations, we found that binary_0.5 yielded the highest multiclass_auc in the test datasets and exhibited less performance degradation compared to training results across all machine learning algorithms (refer to Figure 2B). Following binary_0.5, ranking, qnorm, rma, and ratios transformations were observed. In terms of machine learning algorithms, we observed that glmnet demonstrated the highest multiclass_auc in the test datasets across all data transformations and showed less performance degradation relative to the training phase (see Figure 2C). Succeeding glmnet, rf, svmlinear2, gbm and svmRadial were observed.

#### 3.1.3 Models Interpretation

We calculated the importance of each feature for each model and data transformation combination. The svmLinear2 and svmRadial models utilized all available features (22,277). RandomForest models used between 1,260 and 4,866 features, while gbm models employed fewer features, ranging from 228 to 3,371. The glmnet models used the least features, with counts between 197 and 798 (see Figure 3).

We observed that most selected features were specific to the model and transformation used. For the rma transformation, 72 probes were selected across glmnet, gbm, and rf models. Similarly, 62 probes, 58 probes, 101 probes, and 37 ratios were selected for the qnorm, ranking, binary_0.5, and ratios transformations, respectively (see Supplementary Figure 3A). Within the same model type (glmnet, gbm, rf), there was minimal overlap of probes across different normalizations. No probes overlapped across all five normalizations. The number of common probes for 4 out of 5 transformations was 8 for gbms, 30 for glmnet, and 31 for rf (see Supplementary Figure 3B).

Next, using the probes selected by at least one data transformation for each method, we performed enrichment analysis of biological processes via Gene Ontology (GO) terms (see Supplementary Figure 4), Reactome pathways (see Supplementary Figure 5), KEGG pathways (see Figure 4), and disease ontology semantics (see Figure 4). Our models identified probes whose respective genes are involved in pathways highly related to multiple myeloma, such as the PI3K-Akt, MAPK, JAK-STAT, and Wnt signalling pathways, the RAF/MAP kinase cascade, NF-kB related pathways, and signalling by RAS and BRAF mutants. The disease ontology semantics highlighted several blood cancers, including multiple myeloma, across all methods.

**Figure 4.**
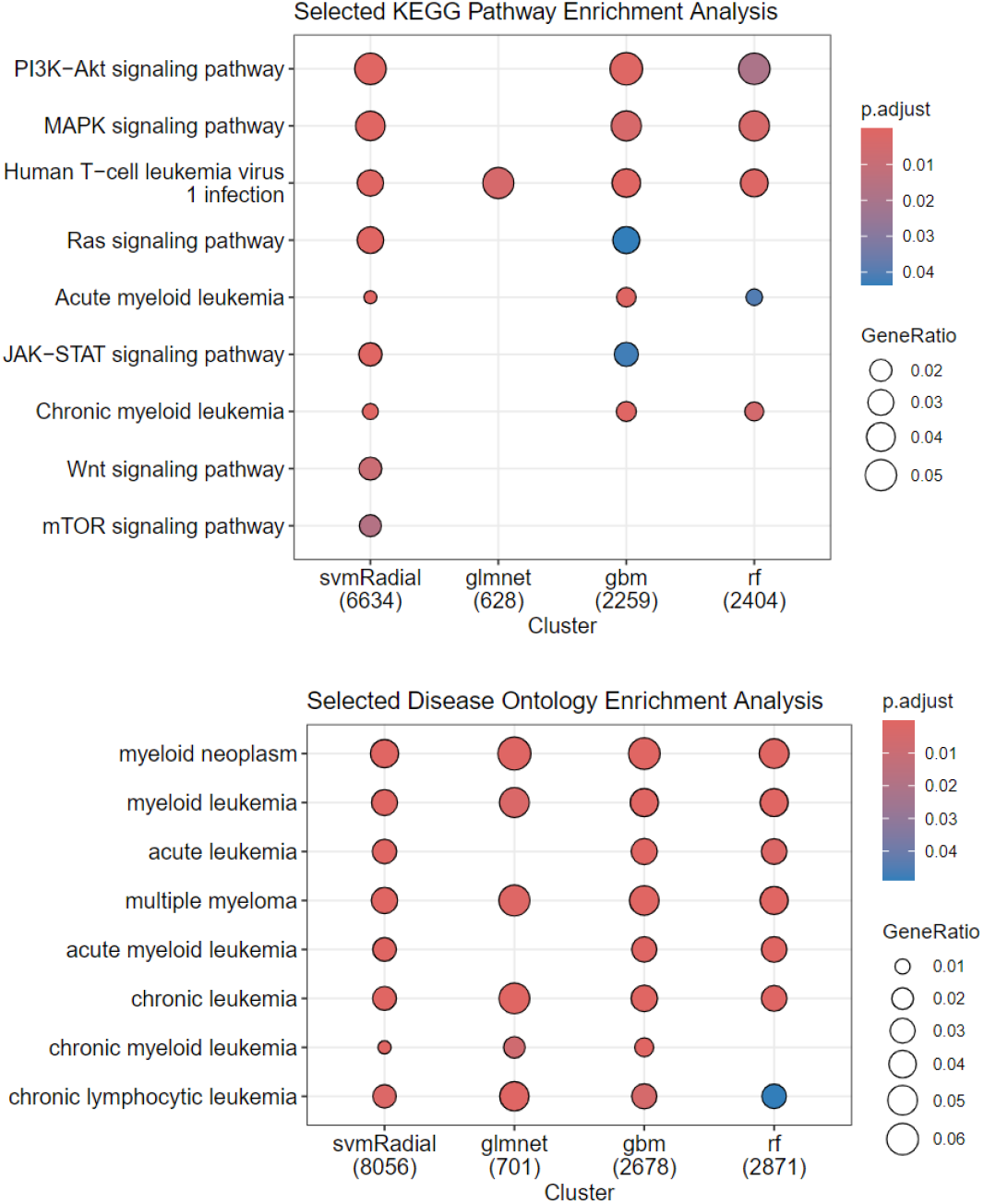
Enrichment analysis for the selected probes. **Top:** KEGG Pathways Associated with Identified Genes. This figure illustrates the KEGG pathways enriched for the genes identified by the machine learning models across different data transformations and training datasets. The pathways displayed are significantly associated with the probes selected by at least one model. Key pathways related to multiple myeloma, such as PI3K-Akt, MAPK, and Wnt signaling, are highlighted. **Bottom:** Disease-related terms associated with Identified Genes. The figure illustrates the distribution of disease-related terms associated with the genes identified by the models. The chart highlights how different methods and data transformations reveal connections to various cancers, including multiple myeloma. Each term represents a disease category. In both figures, the size and color indicate the strength of the association and statistical significance.

**Figure 5.**
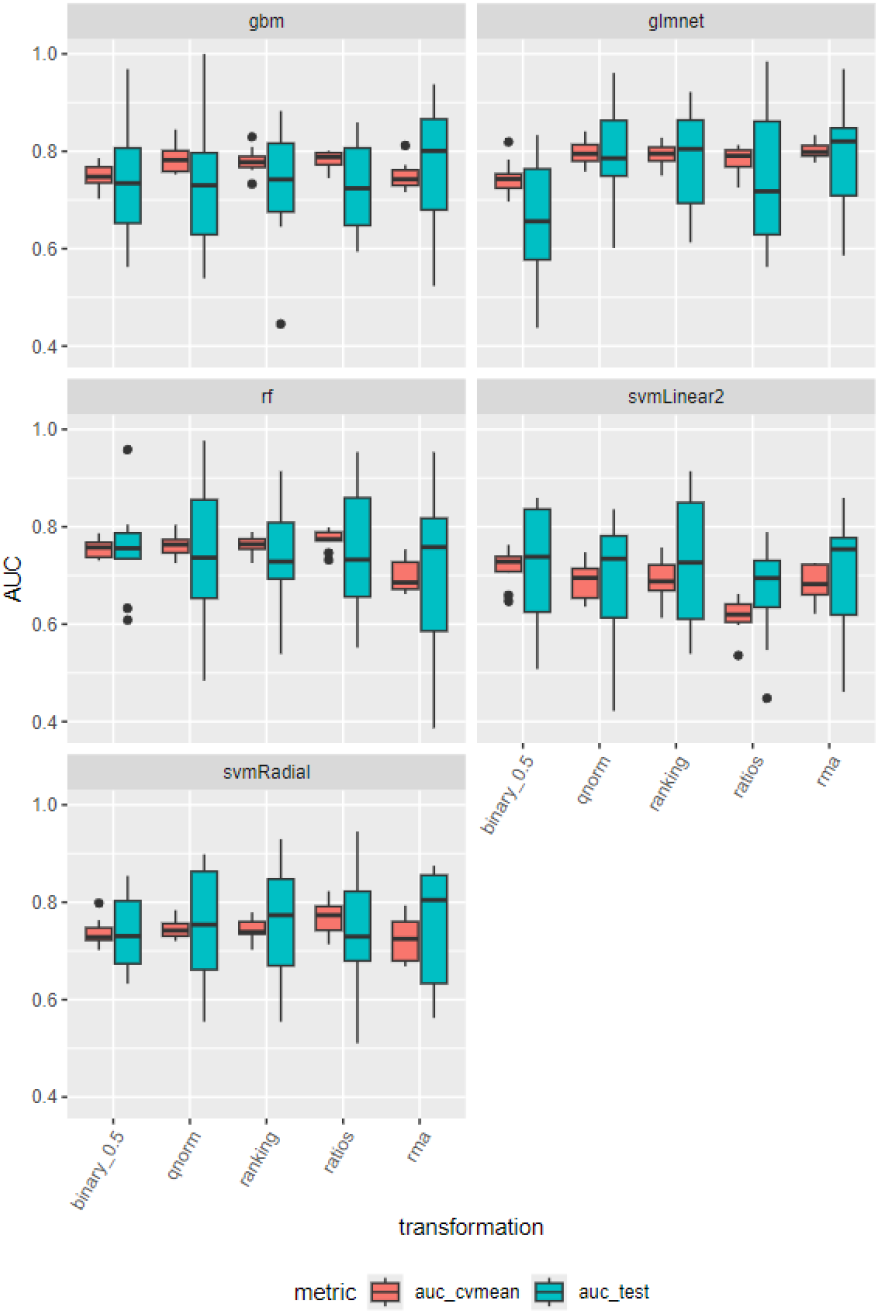
Performance of machine learning algorithms on the GSE235356 dataset. The figure displays the distribution of the mean cross-validation AUC (auc_cvmean, shown in red) and the distribution of the AUC from the outer hold of the nested cross-validation (auc_test, shown in cyan) for each algorithm when the GSE235356 dataset was used for training and testing. The auc_cvmean represents the performance across the cross-validation folds, while the auc_test indicates the model’s generalizability on unseen data. The comparison of these distributions highlights the algorithm’s generalization and stability.

### 3.2 - Task 2. Predicting Progression from MGUS to MM

#### 3.2.1 Model Development for Disease Progression Prediction

We trained our models employing the following datasets: GSE235356, GSE6477, EMTAB317 alone and all datasets generated from the GLP96 and A-AFFY-33 platforms: GSE6477, GSE2113, EMTAB316 and GSE13591. For GSE235356^27^, we trained our models to distinguish between *MGUS and progressing MGUS*, which refers to MGUS cases that progressed to MM. We utilized a 10-fold cross-validation to optimize our model’s hyperparameters and employed a 10-fold nested cross-validation protocol to assess performance. In other scenarios, we trained our models to differentiate between *MGUS and MM*. This involved optimizing the hyperparameters using a ten-fold cross-validation repeated ten times. For consistency, we applied the same machine learning models and data transformations as in task 1, see Figure 1.

The scope of these two training approaches was twofold. First, we aimed to assess whether models trained to differentiate MGUS from MM could effectively distinguish MGUS from progressing MGUS, using the GSE235356 dataset for testing. Second, we sought to compare the performance of these models with those specifically trained to separate MGUS from progressing MGUS patients. This comparison would provide insights into whether models generalized well across related conditions or if specialized training was required for optimal performance in predicting MGUS progression.

#### 3.2.2 Evaluation of Model Performance

Using the ***GSE235356*** dataset for training, we calculated the cross-validation Area Under the ROC Curve (AUC) during model optimization (auc_cv), the mean cross-validation AUC (auc_cvmean), and the outer cross-validation AUC from the nested cross-validation protocol (auc_test). Among the models, glmnet achieved the best performance, followed by gbm, rf, svmRadial, and svmLinear2 (refer to Figure 5). Specifically, glmnet with rma, qnorm, or ranking transformations showed the highest performance, with both auc_cvmean and auc_test around 0.8 (refer to Figure 5). All algorithms and data transformations also demonstrated good generalization performance in the outer cross-validation fold. The mean auc_test across all outer cross-validation folds fell within the AUC distribution achieved during training cross-validation (refer to Figure 6).

**Figure 6.**
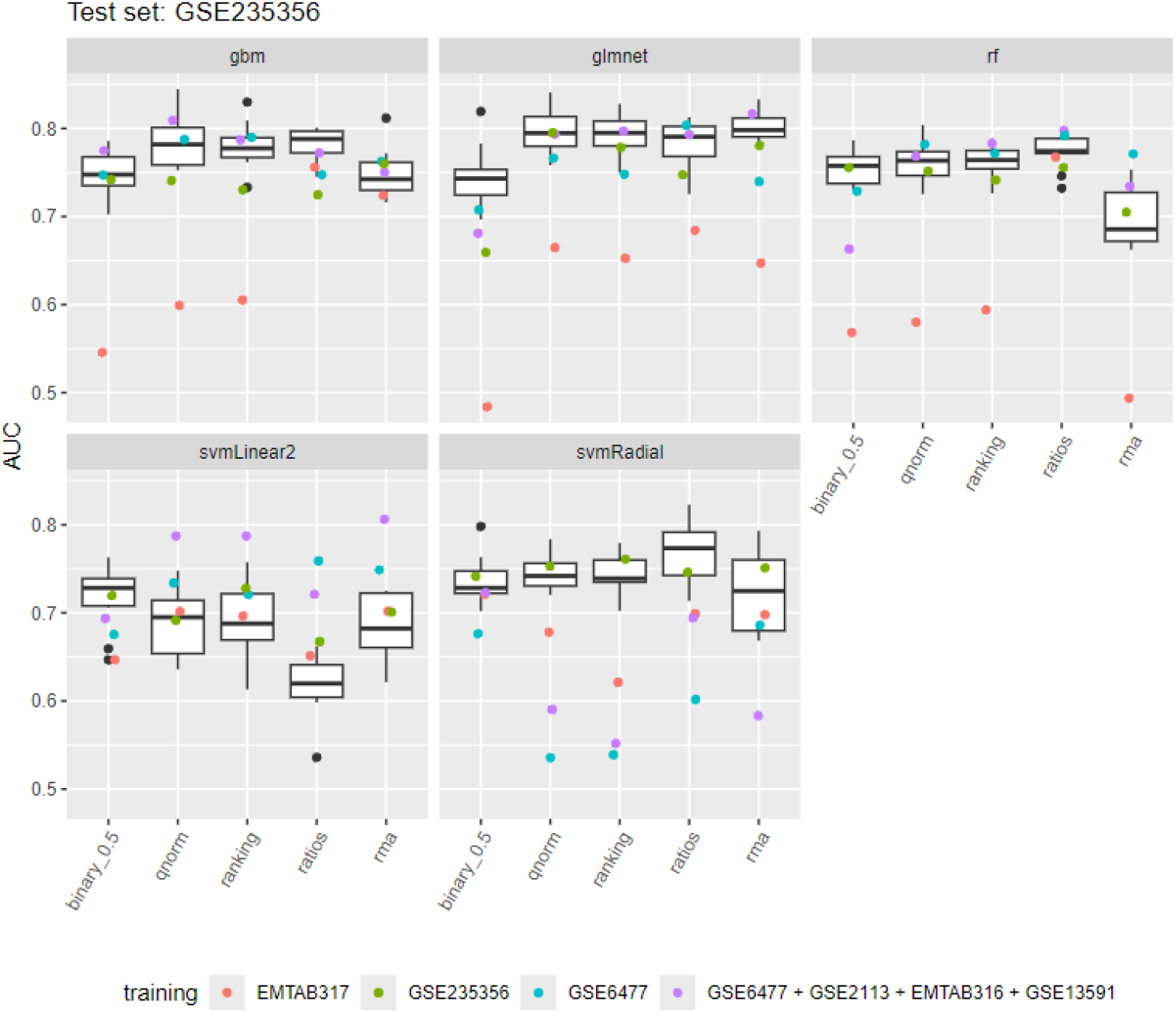
Model performance in differentiating MGUS from progressing MGUS across different datasets. The boxplots show the distribution of the mean cross-validation AUC for models trained to differentiate MGUS from progressing MGUS using the GSE235356 dataset. The colored points represent the performance of each algorithm-data transformation combination across various training datasets: models trained with the EMTAB317 dataset are shown in red; those trained with the GSE235356 dataset are in green; models trained with the GSE6477 dataset are shown in cyan; and those trained with the combined GSE6477 + GSE2113 + EMTAB316 + GSE13591 datasets are depicted in purple. Notably, in all cases except for the second (GSE235356), the models were specifically trained to distinguish MGUS from MM.

We trained our models to distinguish *MGUS from MM* using the ***GSE6477*** dataset. These models achieved a training cross-validation AUC median ranging from 0.93 to 1 across all data transformations and machine-learning methods (refer to Supplementary Figure 6). Most models demonstrated good generalization performance when applied to other datasets (EMTAB316, EMTAB317, GSE13591, GSE2113) for identifying *MGUS from MM* (refer to Supplementary Figure 7). For EMTAB316 and GSE2113, the test AUC median was 0.9 and 0.86 across all data transformations and machine learning methods. For GSE13591, the test AUC median was 0.8 across all methods and transformations. The models achieved the lowest test AUC for the EMTAB317 dataset, with an AUC median of 0.7. This result is consistent with our findings in task 1 and likely due to the different microarray platforms used to generate the data.

**Figure 7.**
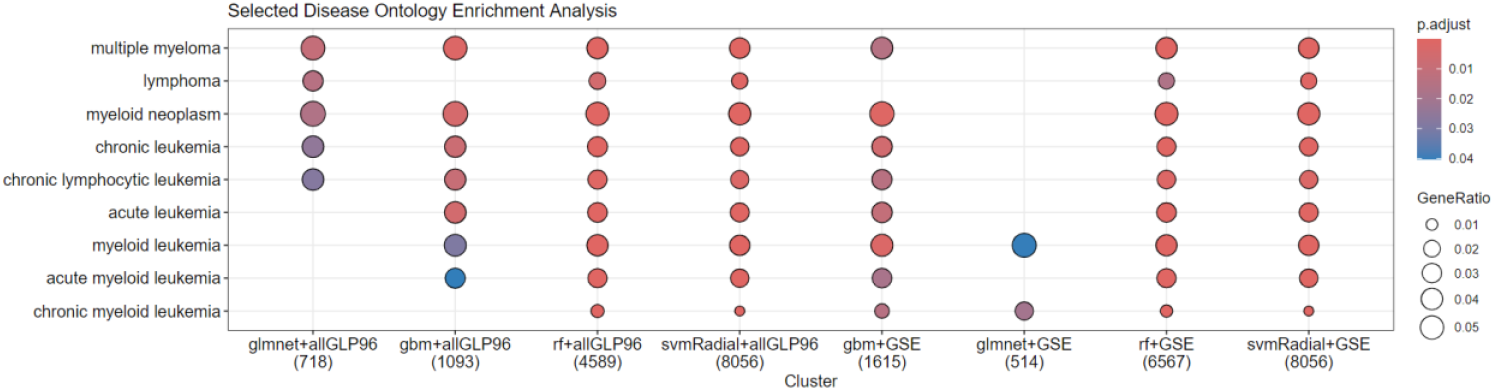
Disease-related terms associated with Identified Genes. The figure illustrates the distribution of disease-related terms associated with the genes identified by the models. The chart highlights how different methods across all data transformations and the different training datasets reveal connections to various cancers, including multiple myeloma. Each term represents a disease category. The size and color indicate the strength of the association and statistical significance. “all GLP96” refers to the combined dataset of GSE6477 + GSE2113 + EMTAB316 + GSE13591, and “GSE” to the GSE235356 dataset.

When we applied our models to separate *MGUS from progressing MGUS*, the models differentiated the two classes. Specifically, the test AUC achieved by gbm, glmnet, rf, and svmLinear2 ranged from 0.7 to 0.8, falling within the AUC distribution achieved with cross-validation during training with the GSE235356 dataset (refer to Figure 6) and within the outer cross-validation auc_test distribution of the GSE235356 dataset (refer to Supplementary Figure 11). In the case of svmRadial, the auc_test ranged from 0.54 to 0.69; in all cases except rma normalization, it was below the training AUC cross-validation distribution but inside the outer cross-validation auc_test distribution of the GSE235356 dataset.

Next, we trained our models to distinguish *MGUS from MM*, employing the ***EMTAB317*** dataset. These models achieved a training cross-validation AUC median ranging from 0.79 to 0.94 across all data transformations and machine-learning methods (refer to Supplementary Figure 8). Most models demonstrated good generalization performance when applied to other datasets (EMTAB316, GSE13591, GSE2113, GSE6477) for identifying *MGUS from MM* (refer to Supplementary Figure 9). For EMTAB316 and GSE6477, the median test AUC was close to 0.75 and 0.82 across all data transformations and machine learning methods. For GSE13591 and GSE2113, the median test AUC was close to 0.86 and 0.91 across all methods and transformations.

When we applied our models to separate *MGUS from progressing MGUS*, the models showed a test AUC performance ranging from 0.5 to 0.76, with a median of 0.65. The test AUC achieved by gbm, glmnet, rf, and svmRadial fell below the cross-validation AUC distribution achieved during training with the GSE235356 dataset (refer to Figure 6) but within the outer cross-validation auc_test distribution of the GSE235356 dataset (refer to Supplementary Figure 11). Interestingly, for svmLinear2, the test AUC fell within the AUC cross-validation distribution during training with the GSE235356 dataset for all data transformations.

Last, we trained our models to separate *MGUS from MM* using all datasets generated from the GLP96 or A-AFFY-33 platforms (***GSE6477 + GSE2113 + EMTAB316 + GSE13591***). Our models achieved a training cross-validation AUC median ranging from 0.94 to 0.97 across all data transformations and machine-learning methods (refer to Supplementary Figure 10). Similarly, we applied our models to separate *MGUS from progressing MGUS*. The models’ test AUC performance ranged from 0.55 for svmRadial using ranking to 0.82 for glmnet employing rma, with a median performance across all models and data transformations of 0.77. Importantly, the test AUC achieved by gbm, glmnet, and rf fell within the cross-validation AUC distribution achieved during training with the GSE235356 dataset (refer to Figure 6) and within the outer cross-validation auc_test distribution of the GSE235356 dataset (refer to Supplementary Figure 11). For svmRadial, the test AUCs fell below the cross-validation AUC distribution for all data transformations except binary_0.5, and at the lower end of the outer cross-validation auc_test distribution. Interestingly, for svmLinear2, the test AUC fell above the cross-validation AUC distribution for all data transformations except binary_0.5, and on the upper end of the outer cross-validation auc_test distribution. Additionally, with the inclusion of the GSE2113, EMTAB316, and GSE13591 datasets, glmnet and svmLinear2 showed a 0.05 increase in median test AUC across all data transformations compared to when only the GSE6477 was used for training; however, these differences were not statistically significant.

Next, we conducted a permutation test to assess the statistical significance of the observed model performances in the test dataset in comparison to a random classification. In this analysis, we permuted the class labels (*MGUS, progressing MGUS*) and recalculated the auc_test for each model. Models with auc_test values close to 0.5, which correspond to a random classifier, did not demonstrate statistically significant different performance from random, as exprected. Conversely, models with auc_test values exceeding 0.7 showed highly significant results, clearly falling outside the permutation distribution (refer to Supplementary 12).

#### 3.2.3 Models Interpretation

We focused on interpreting the models trained using either the GSE235356 dataset or all GPL96 datasets combined. The svmLinear2 and svmRadial models utilized all available features. When all GPL96 datasets were used for training, the rf models employed between 2,169 and 10,269 features, glmnet selected between 236 and 859 probes, and gbm chose between 214 and 685 probes. In contrast, when the GSE235356 dataset was used for training, the rf models utilized between 3,090 and 11,650 features, glmnet selected between 10 and 792 probes, and gbm chose between 47 and 2,084 probes (see Supplementary Figure 13). We also assessed the overlap of probes selected across the two training datasets. For gbm and glmnet, only a small number of probes (ranging from 1 to 99) were selected in both cases. In contrast, the rf models showed a higher degree of overlap, with common probes ranging from 482 to 5,459 (refer to Supplementary Figure 14).

Using the probes selected by at least one data transformation for each method and training dataset, we conducted enrichment analyses on Gene Ontology (GO) biological processes, KEGG pathways, Reactome pathways, and disease ontology semantics. The analysis revealed that our models identified probes associated with genes involved in pathways closely related to multiple myeloma, such as PI3K-Akt (see Supplementary Figure 17), MAPK (see Supplementary Figure 15, 16 and 17), Wnt signalling (see Supplementary Figure 16), BRAF and RAF1 fusion signalling (see Supplementary Figure 17), and mTOR pathways (see Supplementary Figure 16). The disease ontology analysis also underscored the relevance of several blood cancers, including multiple myeloma, across most methods and training datasets (see Figure 7).

## 4 Discussion

In this study, we utilized advanced machine learning (ML) techniques to tackle two critical challenges in multiple myeloma (MM): *predicting the disease stage* and *predicting disease progression* from monoclonal gammopathy of undetermined significance (MGUS) to MM. Through comprehensive data preprocessing, model training, and evaluation across multiple datasets, we aimed to enhance diagnostic precision and offer valuable prognostic insights for hematologic malignancies.

The first focus of our study was on predicting the stage of MM. Accurate staging is crucial for determining the appropriate treatment strategy and prognosis. We developed models using various ML algorithms, including ElasticNet, Random Forest, Boosting, and Support Vector Machines. These models were trained on a dataset comprising samples from different stages of MM and healthy samples, and their performance was evaluated on external validation datasets. The multiclass area under the curve values obtained during cross-validation and testing consistently demonstrated that the selected features and ML algorithms effectively capture the biological differences across disease stages. Among the models evaluated, gbm achieved the highest performance in training, and glmnet showed minimal degradation across different data transformations and datasets, indicating its robustness and generalizability. Our findings align with the growing body of literature that supports the use of ML in oncology, particularly in hematologic malignancies. Previous studies have shown the effectiveness of ML algorithms in improving diagnostic accuracy and risk stratification in MM ^26,38^. The variability in model performance across different platforms, observed in datasets from GPL96, A-AFFY-34, GPL570, and A-AFFY-44, underscores the challenges of integrating data from diverse sources. This issue has been documented in the literature, where differences in data generation methods significantly affect model performance ^39,40^.

Predicting the progression of monoclonal gammopathy of undetermined significance (MGUS) to multiple myeloma (MM) remains one of the most pressing challenges in managing plasma cell disorders. Early identification of high-risk MGUS patients could significantly enhance clinical outcomes by enabling timely interventions that might delay or even prevent the onset of MM. A significant obstacle in this effort is the limited availability of datasets that include progressing MGUS patients, as these cases are inherently rare and difficult to procure. To address this challenge, we employed a two-pronged approaches. First, we developed machine-learning models using a dataset specifically containing *MGUS and progressing MGUS* patients, achieving a maximum AUC of 0.8 with the glmnet model combined with quantile normalization. Other models and data transformations demonstrated good generalization performance, with AUC values around 0.75. This result highlights the potential of machine learning in identifying high-risk MGUS patients even with limited data availability. Second, to evade the scarcity of progressing MGUS samples, we trained our models using multiple datasets containing both *MGUS and MM* patients. These models were then evaluated for their ability to distinguish *MGUS from progressing MGUS* cases. Our findings indicate that machine learning models, including ElasticNet, Boosting, SVM with linear kernel, and Random Forest, achieved AUC values close to 0.8, suggesting a strong potential for these models in risk stratification. Although some models, such as SVM with radial kernel, demonstrated lower performance, the overall results underscore the utility of incorporating both MGUS and MM data in predictive modeling.

To our knowledge, this study is the first to develop comprehensive machine-learning models specifically designed to predict the progression of MGUS to MM by leveraging datasets from both MGUS and MM cases. Our innovative approach of integrating MM data to train models that predict MGUS progression offers a novel and potentially more accurate method for risk assessment. This methodology could have significant clinical implications, particularly in distinguishing MGUS patients who require closer monitoring from those who may not. The novelty and potential impact of our approach are further emphasized by recent reviews in the field, such as the one by Awada et al. ^41^, which highlight the need for more sophisticated predictive models that integrate data across disease stages to enhance prognostication and treatment planning.

The feature selection and enrichment analyses conducted in this study provided significant insights into the molecular pathways and biological processes involved in the progression of multiple myeloma. Our models consistently identified genes involved in critical signaling pathways, such as PI3K-Akt, MAPK, Wnt, and mTOR. These pathways are well-known for their roles in cell growth, survival, and proliferation, and their involvement in MM pathogenesis is well-documented ^42^. For instance, the PI3K-Akt pathway has been widely recognized as a key player in MM, influencing proliferation, migration, apoptosis, and autophagy ^43^. Similarly, the MAPK pathway is involved in the regulation of cell proliferation, survival, and differentiation, and its dysregulation has been implicated in various cancers, including MM ^44,45^. The Wnt pathway, which is crucial for cell differentiation and proliferation, has also been associated with MM progression, particularly in the context of bone disease ^46^. The consistency of our results with established biological knowledge validates our models and suggests potential therapeutic targets that could be explored in future research.

While the results of this study are promising, several limitations should be considered when interpreting our findings. One significant challenge is the variability in model performance across different microarray platforms. This variability suggests a need for more comprehensive cross-platform validation to ensure the robustness of our models in different clinical settings. Moreover, the relatively small number of datasets used in this study and the focus on a limited set of machine learning algorithms may have constrained our ability to explore other potentially valuable approaches. Future research should aim to include a more extensive variety of datasets, especially those generated from different omics technologies, to enhance the generalizability and robustness of the models. Also, to improve model robustness, an ensemble classification approach could be explore. By combining multiple machine learning algorithms, ensemble methods can reduce performance variability across platforms and enhance prediction accuracy, offering more reliable generalizability for clinical use. Additionally, integrating clinical data, such as patient demographics and treatment history, could provide a more comprehensive understanding of disease progression and improve the clinical applicability of the models. These challenges are well-recognized in the literature ^47–49^. All studies emphasize the need for cross-platform validation and standardization in ML models, particularly in the context of precision medicine, where the ability to generalize across different datasets is crucial for clinical implementation.

## 5 Funding

Funded by the European Union. Views and opinions expressed are however those of the author(s) only and do not necessarily reflect those of the European Union or ELMUMY (Project: 101097094). Neither the European Union nor the granting authority can be held responsible for them.

## Notes

### Competing Interest Statement

The authors have declared no competing interest.

