## Supplementary for "Machine Learning Models for Predicting Multiple Myeloma Staging and MGUS Progression Using Gene Expression Data"

#### Task 1: Predicting the Stage of Multiple Myeloma

##### Performance during cross-validation

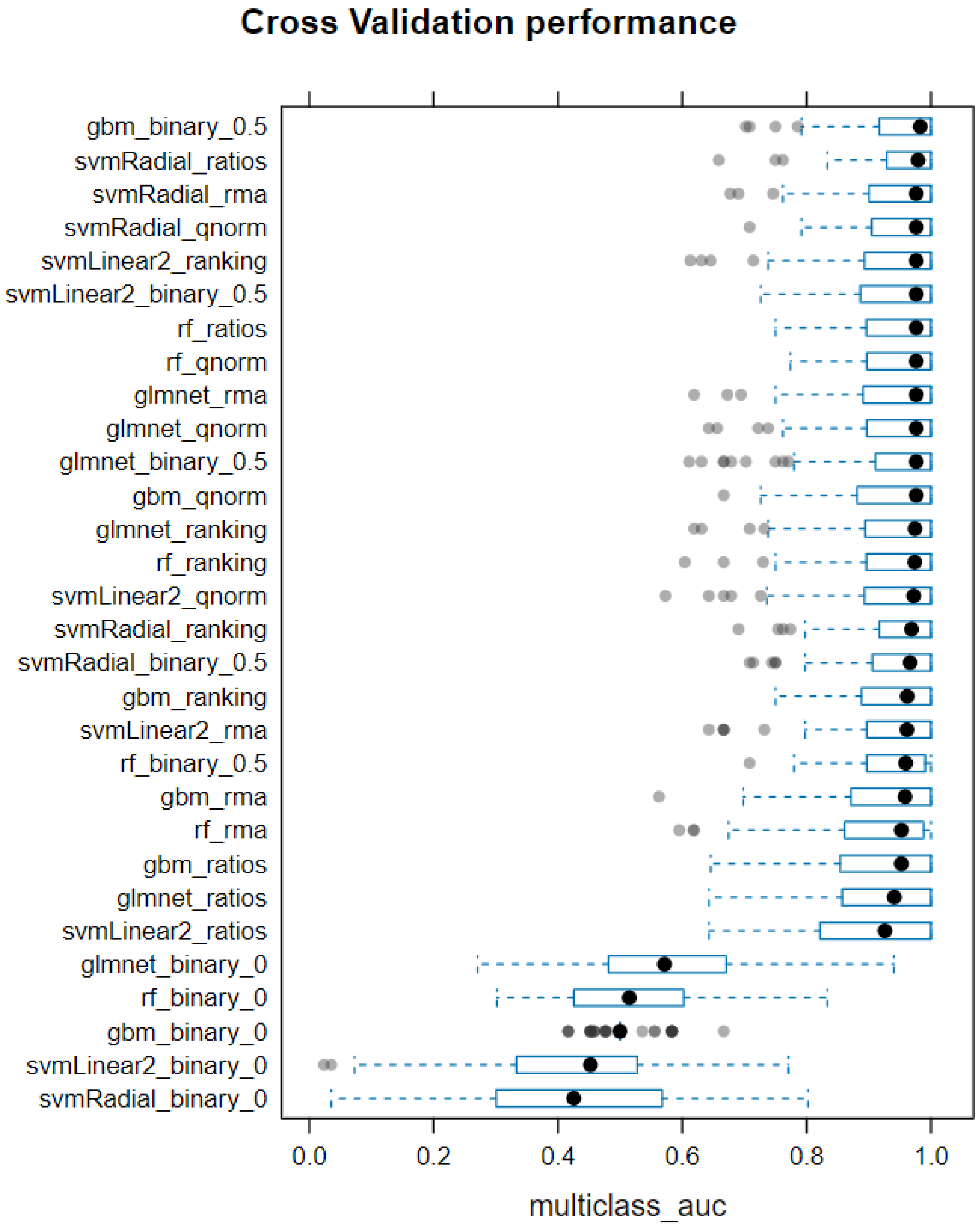


***Supplementary Figure 1. Distribution of Multiclass AUC from Ten-Fold Repeated Cross-Validation.*** *The figure shows the distribution of the multiclass AUC metric across ten-fold cross-validation repeated ten times during training. Models are arranged in descending order, with the best-performing models positioned at the top.*

#### Performance in test datasets

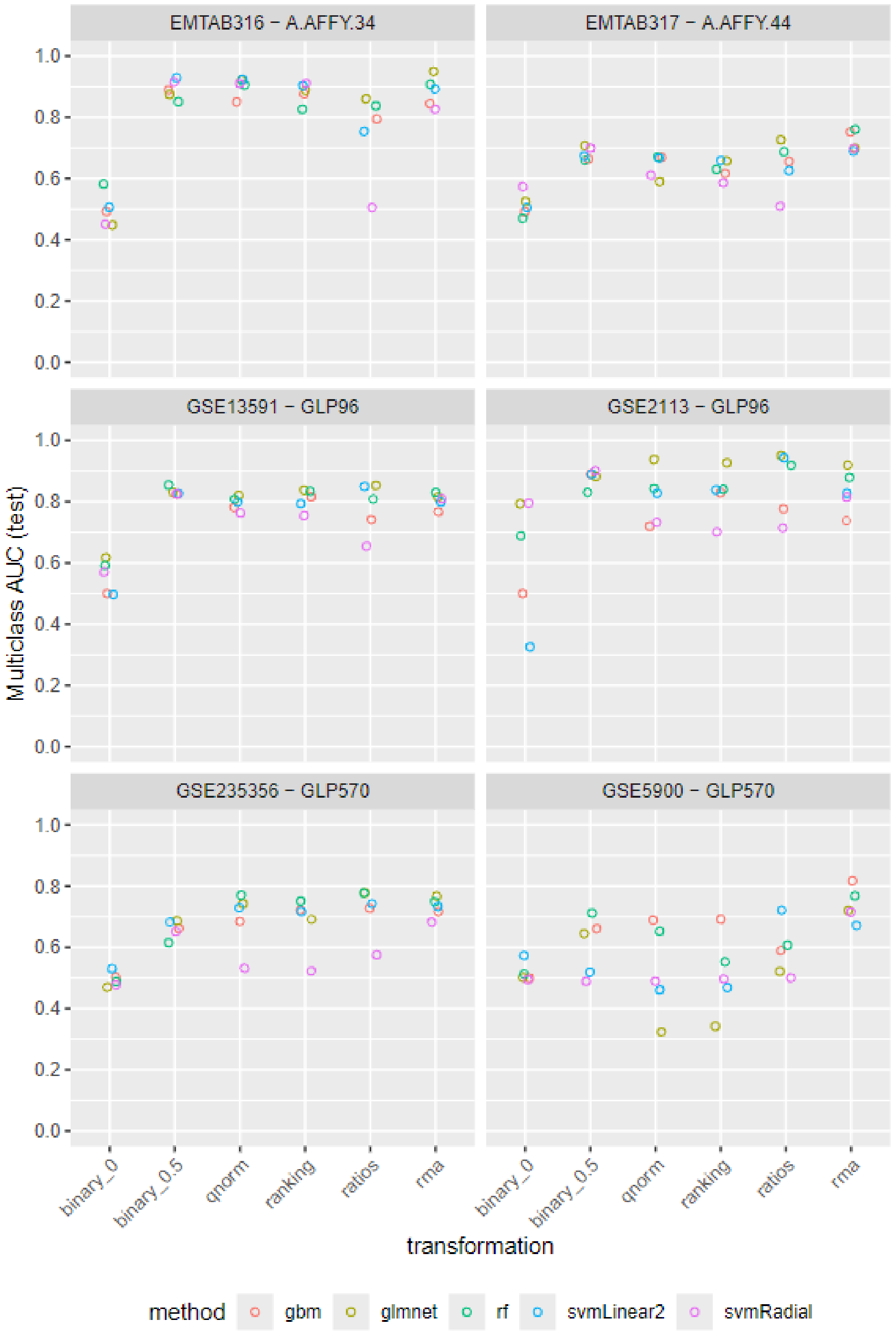


***Supplementary Figure 2. Performance of Machine Learning Model-Data Transformation Combinations on External Datasets.*** *The figure illustrates the performance of each machine learning model with different data transformations across external datasets. The performance of gbm is represented in red, glmnet in yellow, rf in green, svmLinear2 in blue, and svmRadial in purple.*

#### Model interpretation

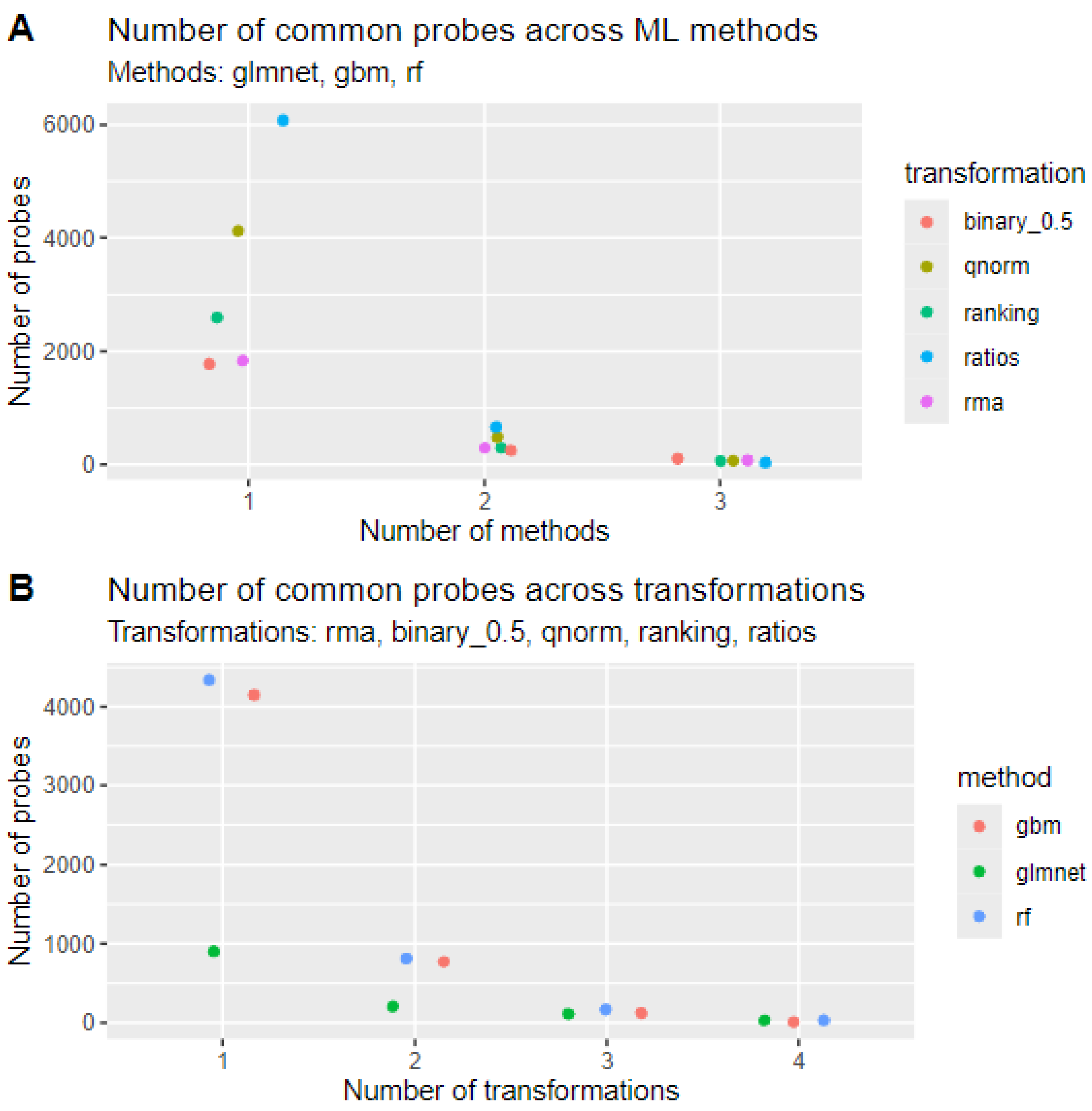


***Supplementary Figure 3. Number of Common Probes Selected.*** *A) The number of common probes selected across different machine learning methods (glmnet, gbm, and rf) for each data transformation. B) The number of common probes selected across various data transformations for each machine learning method.*

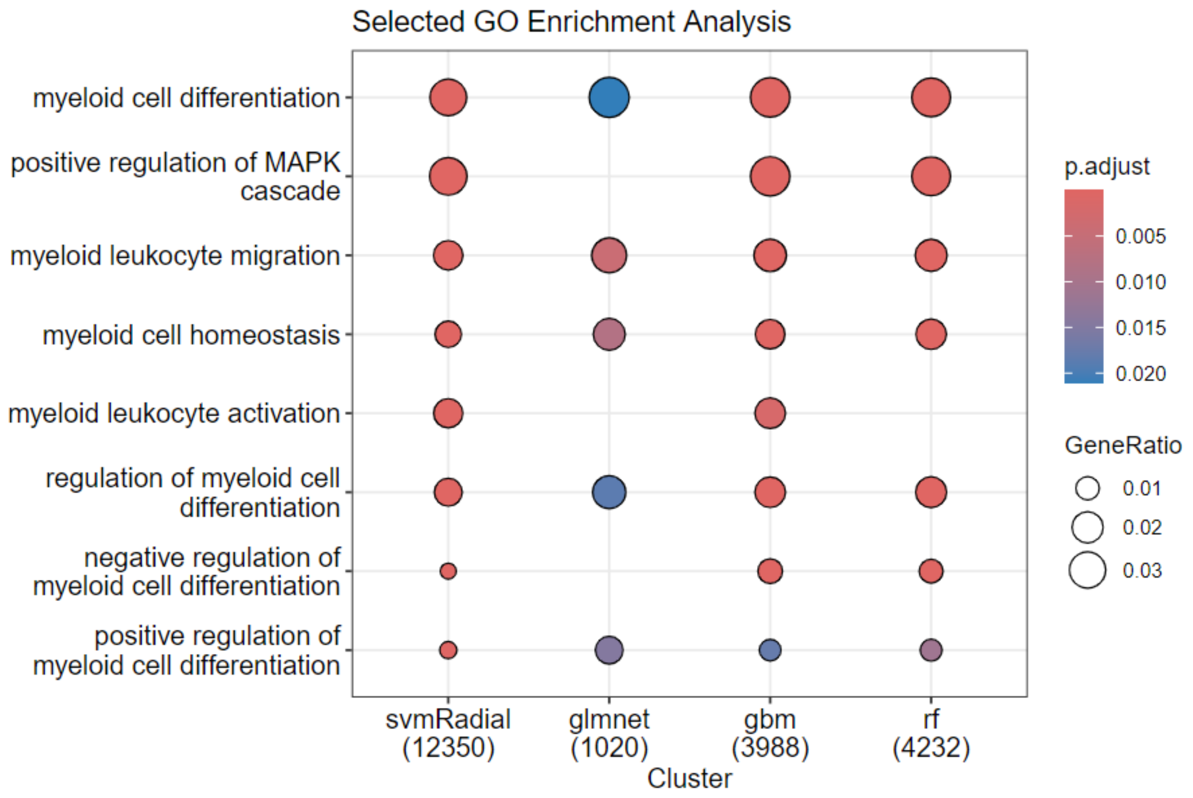


***Supplementary Figure 4. GO Enrichment Analysis of Identified Genes.*** *This figure presents the Gene Ontology (GO) enrichment analysis for the genes identified by the machine learning models across all data transformations. The analysis focuses on biological processes, highlighting those that are significantly overrepresented among the selected genes and are related to multiple myeloma based on the current literature. The size and color indicate the strength of the association and statistical significance.*

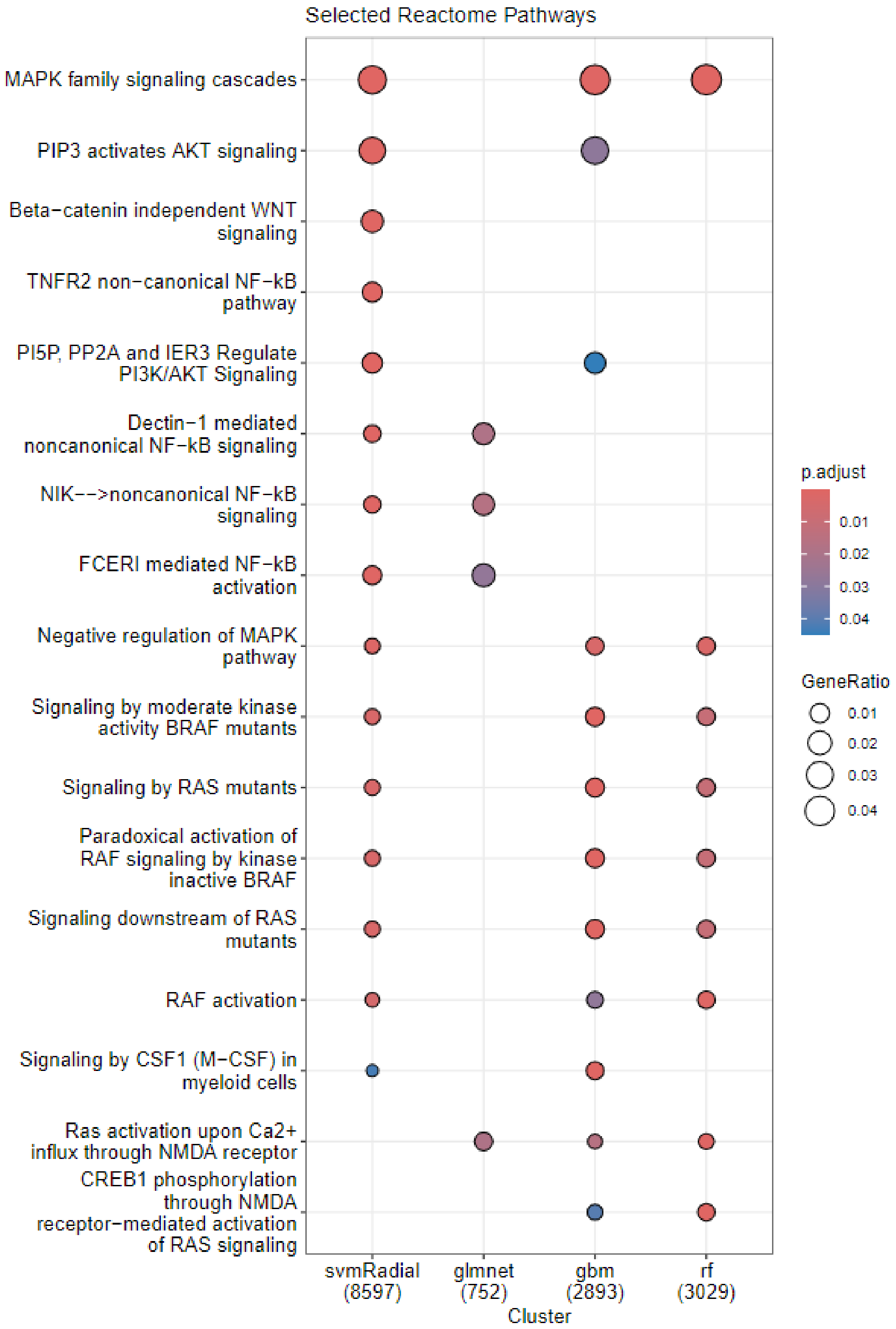


***Supplementary Figure 5. Reactome Pathways Enrichment Analysis of Identified Genes.*** *This figure displays the results of the Reactome Pathways enrichment analysis for genes identified by machine learning models across all data transformations. The highlighted pathways are significantly associated with multiple myeloma and validated by existing literature. The size of the markers indicates the strength of the association, while the color gradient represents the level of statistical significance.*

### Task 2: Predicting Progression from MGUS to MM

#### Training GSE6477

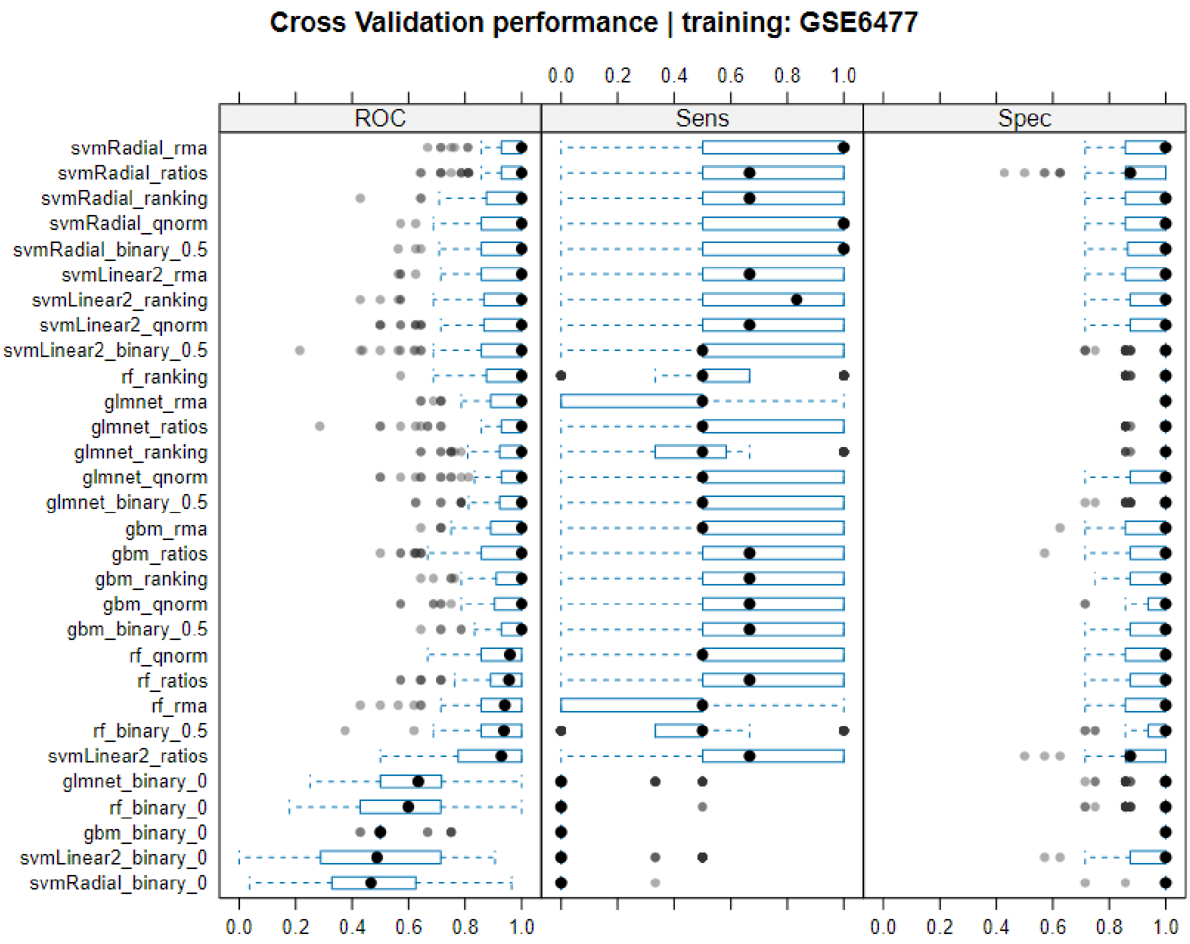


***Supplementary Figure 6. Distribution of AUC from Ten-Fold Repeated Cross-Validation.*** *The figure shows the distribution of the AUC (ROC), Sensitivity (Sens) and Specificity (Spec) metrics across ten-fold cross-validation repeated ten times during training. Models are arranged in descending order, with the best-performing models positioned at the top. The GSE6477 dataset was utilized for training, specifically for distinguishing MGUS from MM.*

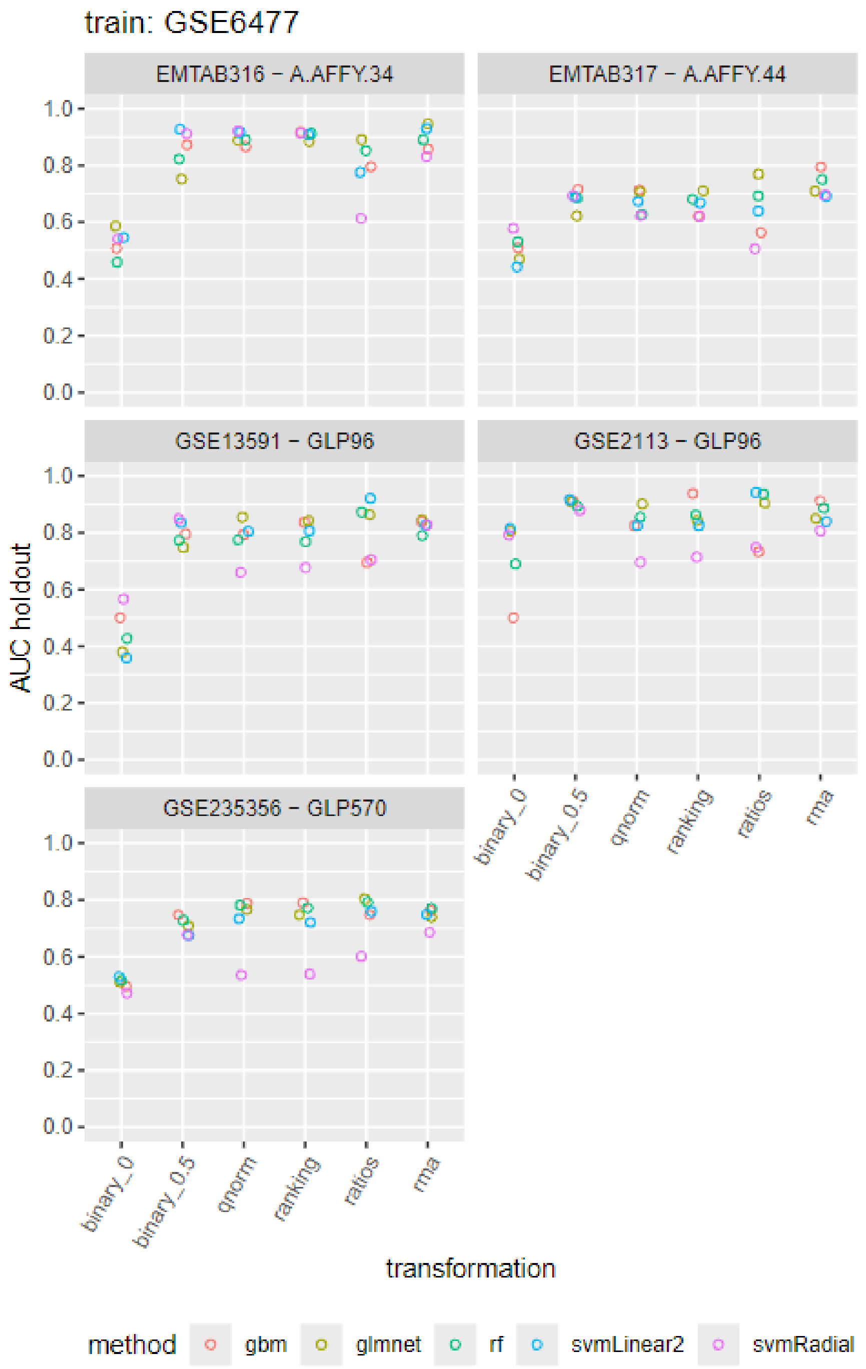


***Supplementary Figure 7. Performance of Machine Learning Model-Data Transformation Combinations on External Datasets.*** *The figure presents the AUC performance of various machine learning models trained on GSE6477, combined with different data transformations across external datasets. For all datasets except GSE235356, the task was to separate MGUS from MM. In the GSE235356 case, they separate MGUS from Progressing MGUS. The performance of gbm is shown in red, glmnet in yellow, rf in green, svmLinear2 in blue, and svmRadial in purple.*

#### Training EMTAB317

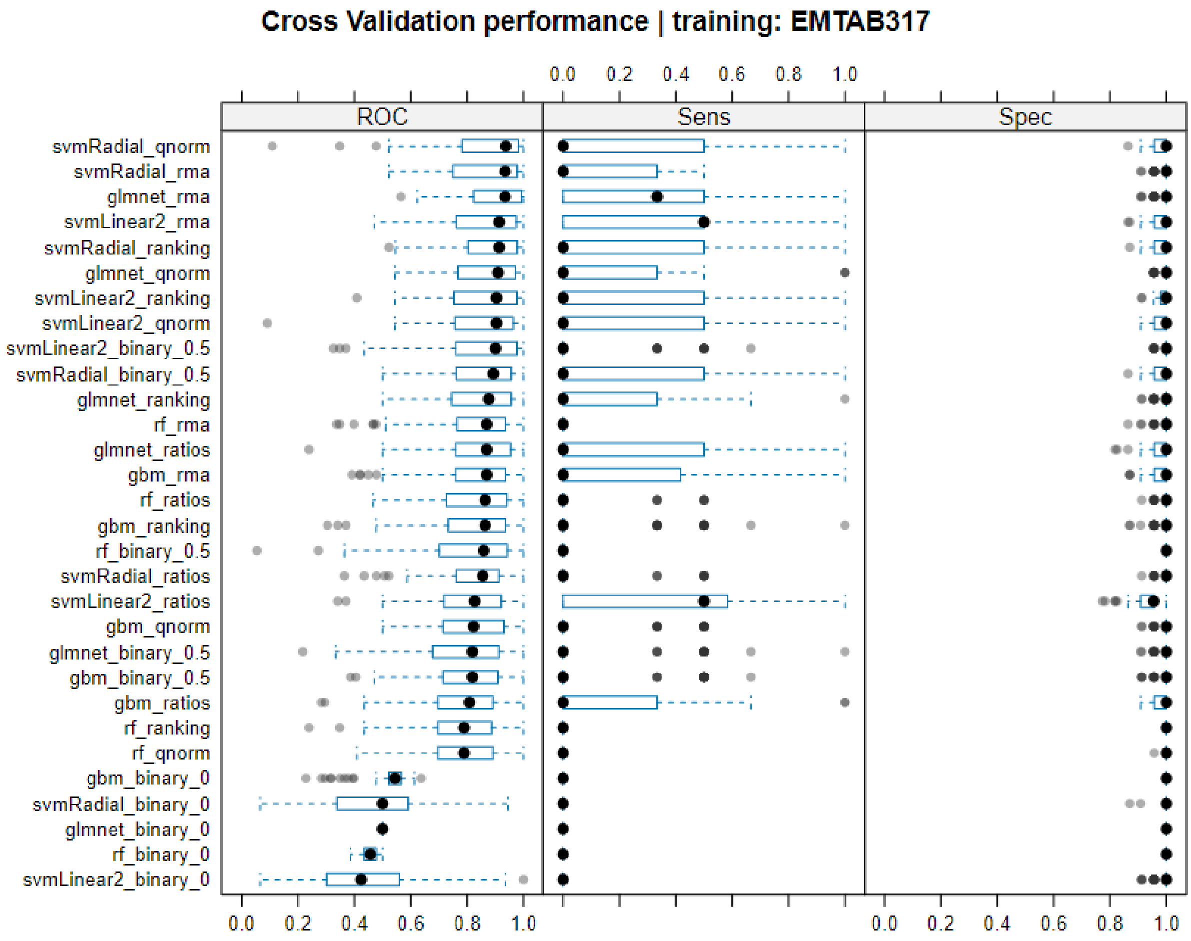


***Supplementary Figure 8. Distribution of AUC from Ten-Fold Repeated Cross-Validation.*** *The figure shows the distribution of the AUC (ROC), Sensitivity (Sens) and Specificity (Spec) metrics across ten-fold cross-validation repeated ten times during training. Models are arranged in descending order, with the best-performing models positioned at the top. The EMTAB317 dataset was utilized for training, specifically for distinguishing MGUS from MM.*

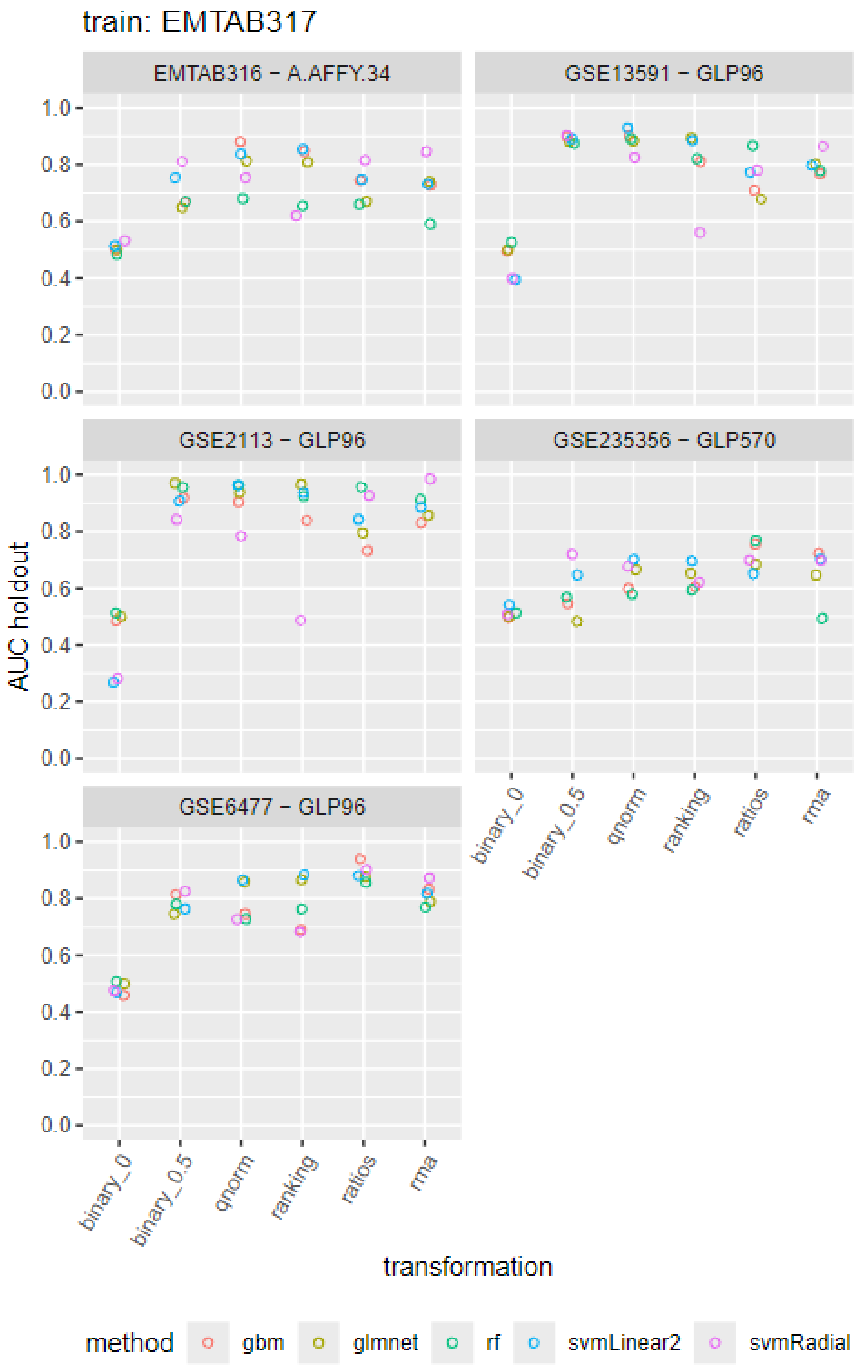


***Supplementary Figure 9. Performance of Machine Learning Model-Data Transformation Combinations on External Datasets.*** *The figure presents the AUC performance of various machine learning models trained on EMTAB317, combined with different data transformations across external datasets. For all datasets except GSE235356, the task was to separate MGUS from MM. In the GSE235356 case, they separate MGUS from Progressing MGUS. The performance of gbm is shown in red, glmnet in yellow, rf in green, svmLinear2 in blue, and svmRadial in purple.*

#### Training GSE6477 + GSE2113 + EMTAB316 + GSE13591

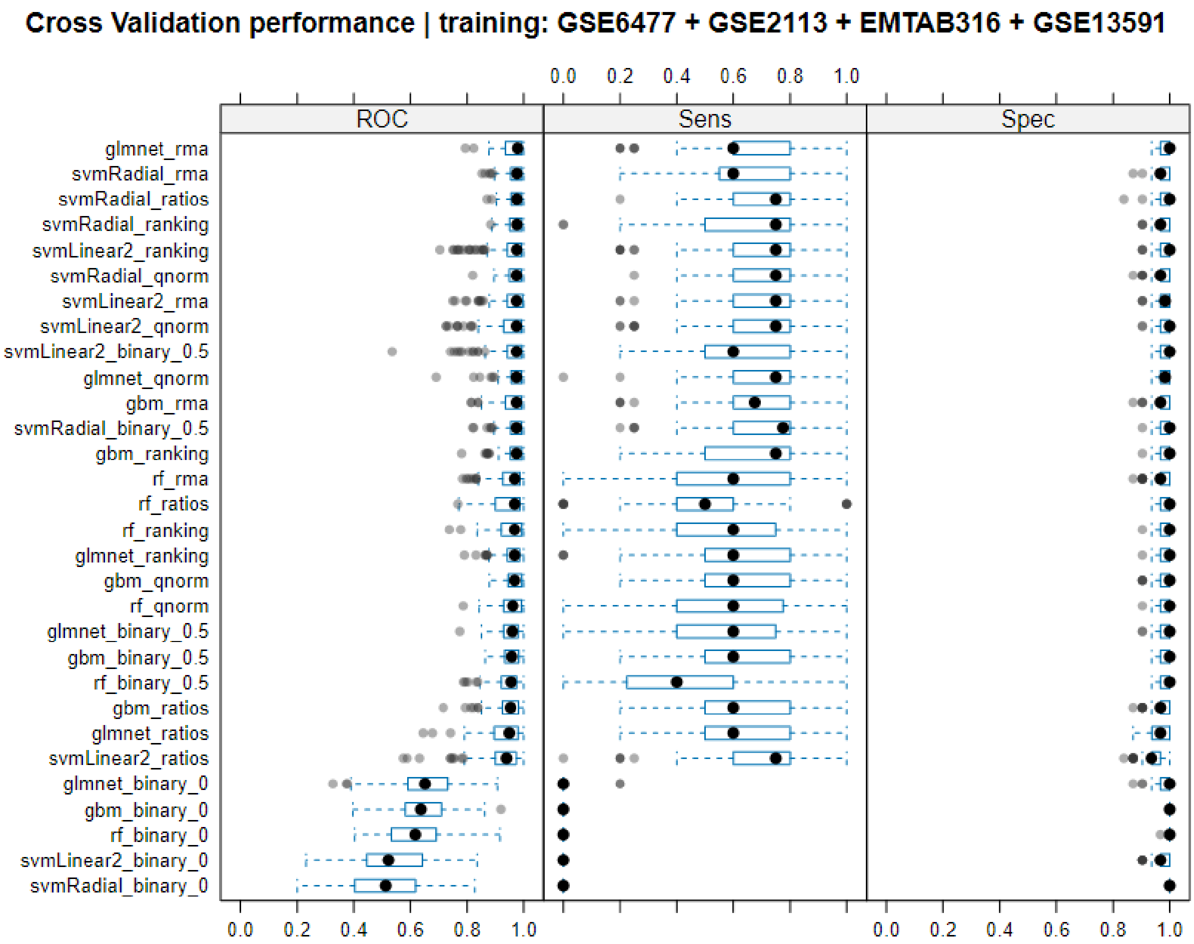


***Supplementary Figure 10. Distribution of AUC from Ten-Fold Repeated Cross-Validation.*** *The figure shows the distribution of the AUC (ROC), Sensitivity (Sens) and Specificity (Spec) metrics across ten-fold cross-validation repeated ten times during training. Models are arranged in descending order, with the best-performing models positioned at the top. The GSE6477 + GSE2113 + EMTAB316 + GSE13591 datasets were utilized for training, specifically for distinguishing MGUS from MM.*

#### Performance on GSE235356.

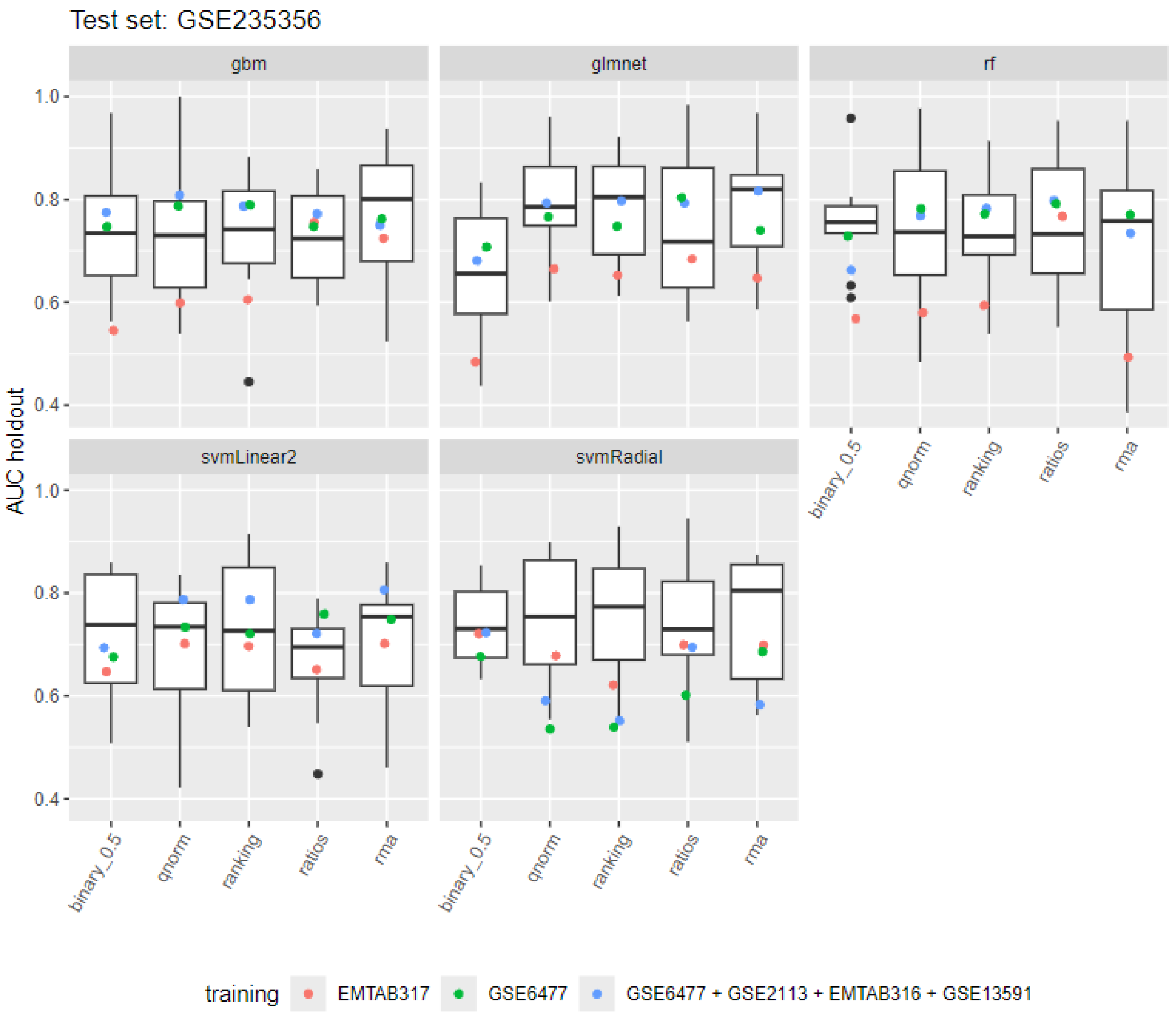


***Supplementary Figure 11. Model performance in differentiating MGUS from progressing MGUS across different datasets.*** *The boxplots show the distribution of the AUC from the outer hold of the nested cross-validation for each algorithm-data transformation combination when the GSE235356 dataset was used for training and testing. The colored points represent the performance of each algorithm-data transformation combination across various training datasets: models trained with the EMTAB317 dataset are shown in red; those trained with the GSE6477 dataset are in green; and those trained with the combined GSE6477 + GSE2113 + EMTAB316 + GSE13591 datasets are depicted in blue. Notably, in all cases, the models were specifically trained to distinguish MGUS from MM.*

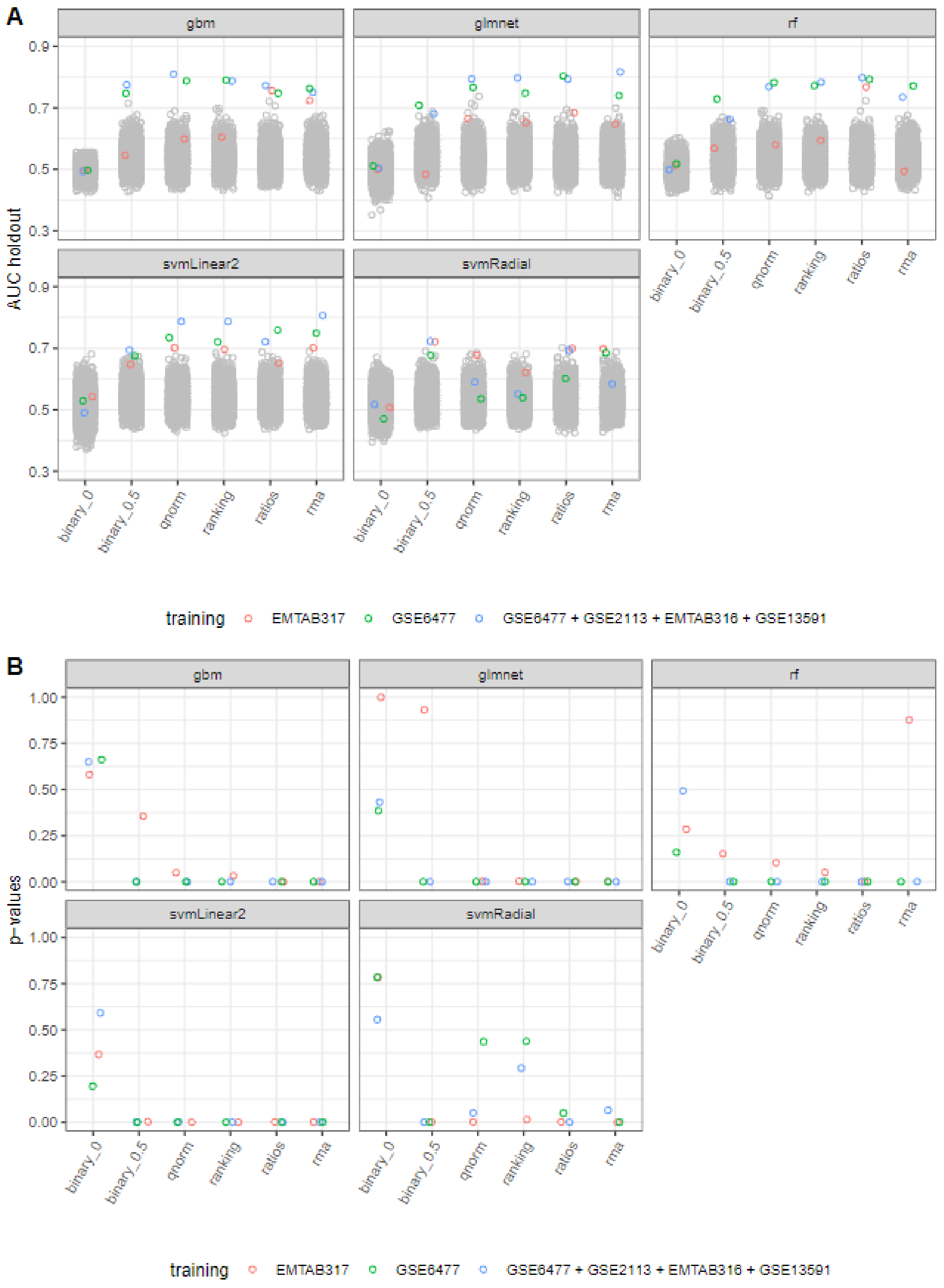


***Supplementary Figure 12. Permutation Testing. A)*** *The figure displays the distribution of permutation-based AUC performance (shown in grey) alongside the AUC performance of each algorithm-data transformation combination across different training datasets. Models trained with the EMTAB317 dataset are represented in red, those trained with the GSE6477 dataset in green, and models trained with the combined GSE6477 + GSE2113 + EMTAB316 + GSE13591 datasets are shown in blue.* ***B)*** *Corresponding permutation-based p-values are provided, illustrating the statistical significance of the observed AUC performances relative to the permutation distribution.*

#### Model Interpretation

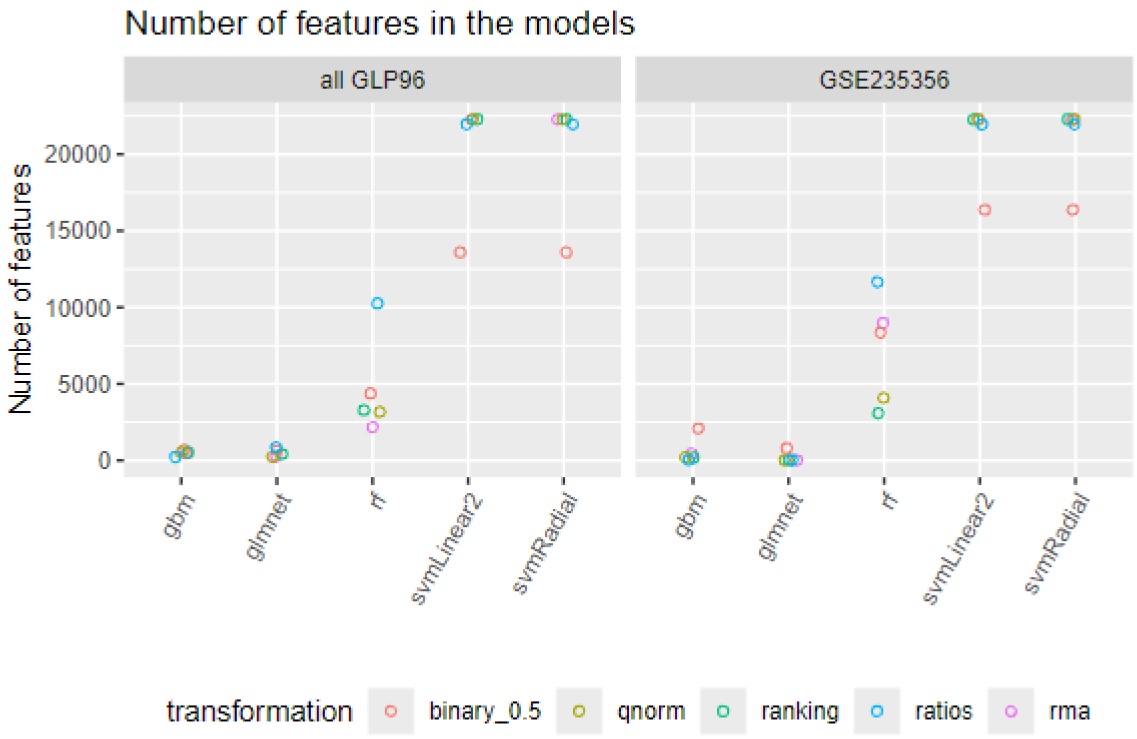


***Supplementary Figure 13. Feature Utilization Across Models and Data Transformations.*** *The figure illustrates the number of features selected by each machine learning model across various data transformations and training datasets. The left panel ("all GLP96") shows the feature selection when models were trained using the combined GSE6477 + GSE2113 + EMTAB316 + GSE13591 datasets, while the right panel displays feature utilization when the GSE235356 dataset was used for training. The plot highlights the variation in the number of features each model utilized, underscoring the differences in feature selection strategies across different datasets and transformations.*

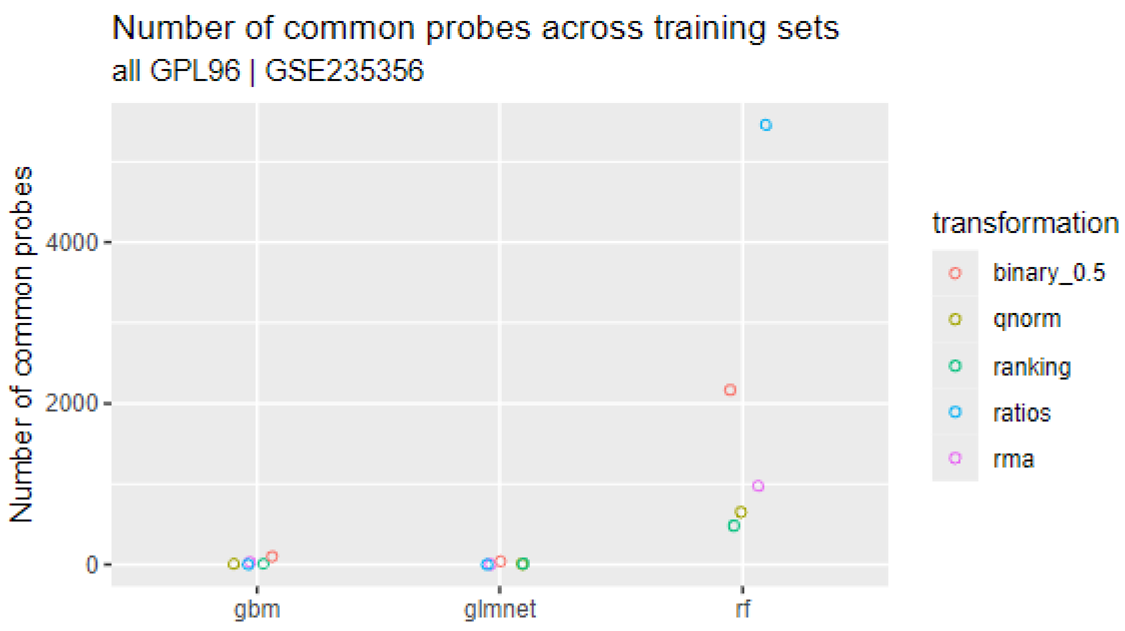


***Supplementary Figure 14. Number of Common Probes Selected Across Training Datasets.*** *The figure displays the number of common probes selected by each machine learning method (glmnet, gbm, and rf) across different data transformations for two training datasets: "all GLP96" and GSE235356. "all GLP96" refers to the combined dataset of GSE6477 + GSE2113 + EMTAB316 + GSE13591.*

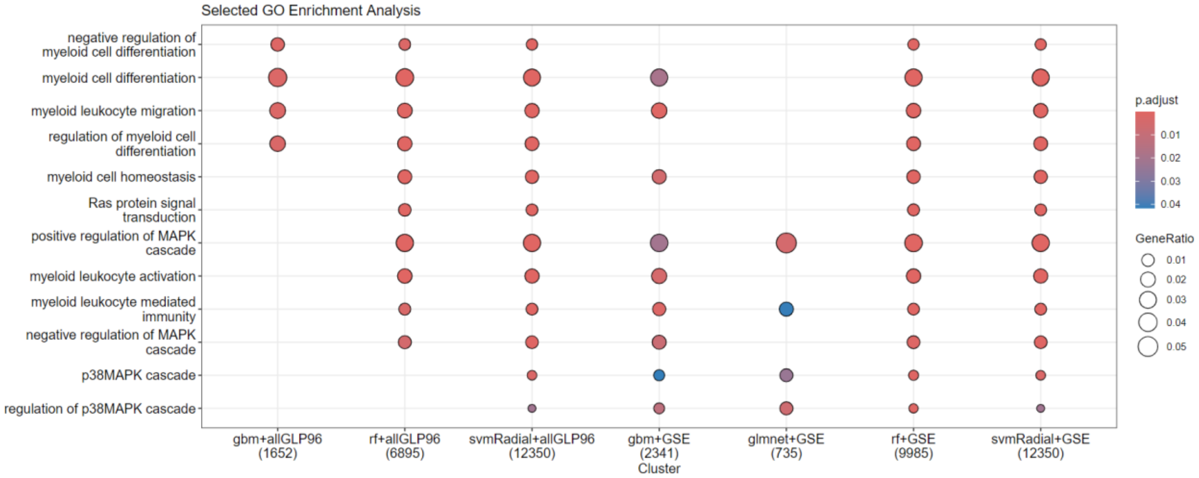


***Supplementary Figure 15. Enrichment Analysis of Selected Gene Ontology (GO) Biological Processes.*** *This figure presents the GO enrichment analysis for genes identified by machine learning models across all data transformations and the different training datasets. The figure focuses on biological processes, emphasizing those significantly overrepresented among the selected genes and closely associated with multiple myeloma according to current literature. Marker size indicates the strength of the association, while the color gradient represents the statistical significance. “all GLP96” refers to the combined dataset of GSE6477 + GSE2113 + EMTAB316 + GSE13591, and “GSE” to the GSE235356 dataset.*

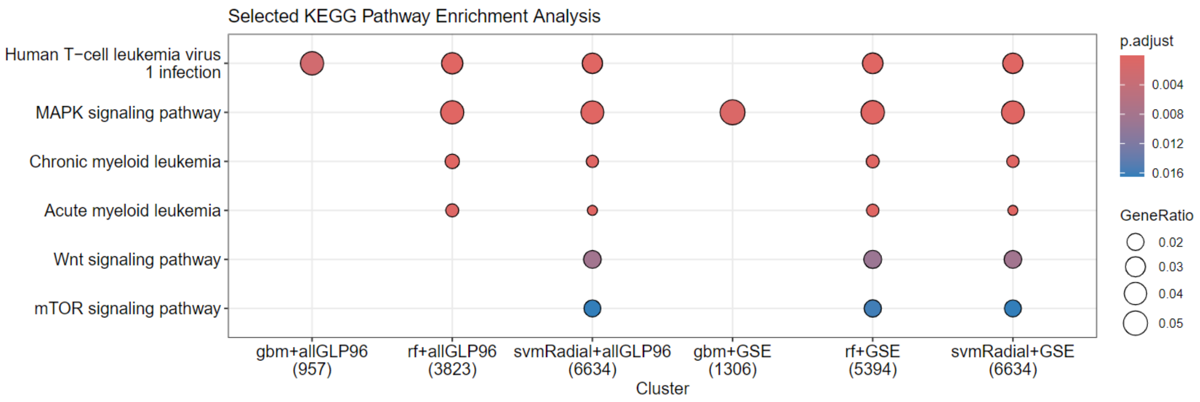


***Supplementary Figure 16. KEGG Pathways Associated with Identified Genes.*** *This figure illustrates the KEGG pathways enriched for the genes identified by the machine learning models across all data transformations and the different training datasets. The pathways displayed are significantly associated with the probes selected by at least one model. Key pathways related to multiple myeloma, such as MAPK, mTOR, and Wnt signaling, are highlighted. "all GLP96" refers to the combined dataset of GSE6477 + GSE2113 + EMTAB316 + GSE13591, and "GSE" to the GSE235356 dataset.*

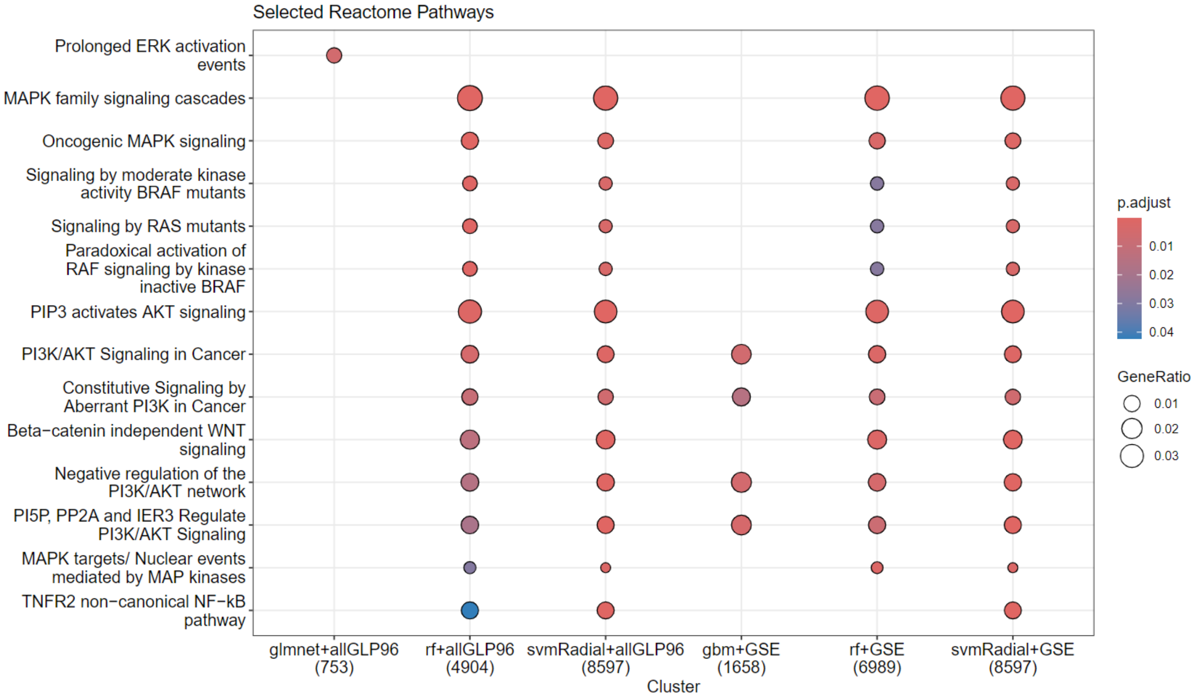


***Supplementary Figure 17. Reactome Pathways Enrichment Analysis of Identified Genes.*** *This figure displays the results of the Reactome Pathways enrichment analysis for genes identified by machine learning models across all data transformations and the different training datasets. The highlighted pathways are significantly associated with multiple myeloma and validated by existing literature. The size of the markers indicates the strength of the association, while the color gradient represents the level of statistical significance. "all GLP96" refers to the combined dataset of GSE6477 + GSE2113 + EMTAB316 + GSE13591, and "GSE" to the GSE235356 dataset.*
